# Characterizing spatiotemporal population receptive fields in human visual cortex with fMRI

**DOI:** 10.1101/2023.05.02.539164

**Authors:** Insub Kim, Eline R. Kupers, Garikoitz Lerma-Usabiaga, Kalanit Grill-Spector

## Abstract

The use of fMRI and computational modeling has advanced understanding of spatial characteristics of population receptive fields (pRFs) in human visual cortex. However, we know relatively little about the spatiotemporal characteristics of pRFs because neurons’ temporal properties are one to two orders of magnitude faster than fMRI BOLD responses. Here, we developed an image-computable framework to estimate spatiotemporal pRFs from fMRI data. First, we developed a simulation software that predicts fMRI responses to a time varying visual input given a spatiotemporal pRF model and solves the model parameters. The simulator revealed that ground-truth spatiotemporal parameters can be accurately recovered at the millisecond resolution from synthesized fMRI responses. Then, using fMRI and a novel stimulus paradigm, we mapped spatiotemporal pRFs in individual voxels across human visual cortex in 10 participants. We find that a compressive spatiotemporal (CST) pRF model better explains fMRI responses than a conventional spatial pRF model across visual areas spanning the dorsal, lateral, and ventral streams. Further, we find three organizational principles of spatiotemporal pRFs: (i) from early to later areas within a visual stream, spatial and temporal integration windows of pRFs progressively increase in size and show greater compressive nonlinearities, (ii) later visual areas show diverging spatial and temporal integration windows across streams, and (iii) within early visual areas (V1-V3), both spatial and temporal integration windows systematically increase with eccentricity. Together, this computational framework and empirical results open exciting new possibilities for modeling and measuring fine-grained spatiotemporal dynamics of neural responses in the human brain using fMRI.

**Significance Statement:** We developed a computational framework for estimating spatiotemporal receptive fields of neural populations using fMRI. This framework pushes the boundary of fMRI measurements, enabling quantitative evaluation of neural spatial and temporal processing windows at the resolution of visual degrees and milliseconds, which was thought to be unattainable with fMRI. We not only replicate well-established visual field and pRF size maps, but also estimates of temporal summation windows from electrophysiology. Notably, we find that spatial and temporal windows as well as compressive nonlinearities progressively increase from early to later visual areas in multiple visual processing streams. Together, this framework opens exciting new possibilities for modeling and measuring fine-grained spatiotemporal dynamics of neural responses in the human brain using fMRI.

## Introduction

The visual scene changes over space and time. To interpret this rich visual input, the visual system processes information spatially and temporally through computations by receptive fields. Prior research has separately characterized spatial receptive fields in primate (Hubel and Wiesel, 1968) and human visual cortex (Dumoulin and Wandell, 2008; Wandell et al., 2009; Kay et al., 2013, 2015; Wandell and Winawer, 2015; Klink et al., 2021) as well as temporal properties of neural responses in primates (Maunsell and Gibson, 1992; Nowak and Bullier, 1997) and humans (Stigliani et al., 2017, 2019; Zhou et al., 2018, 2019; Harvey et al., 2020; Groen et al., 2022; Hendrikx et al., 2022). However, how spatiotemporal information is jointly processed by receptive fields is not well understood beyond primary visual cortex, V1, (McLean and Palmer, 1989; DeAngelis et al., 1993; De Valois and Cottaris, 1998; De Valois et al., 2000; Conway and Livingstone, 2003; Nishimoto et al., 2011) and motion-selective areas, MT/MST, (Simoncelli and Heeger, 1998; Nishimoto and Gallant, 2011; Mineault et al., 2012; Pawar et al., 2019). Thus, it is unknown what are the characteristics of spatiotemporal population receptive fields (pRFs) in human visual cortex.

There are two main reasons for this gap in knowledge. First, measurements of pRFs in humans are derived from fMRI, which typically measures BOLD signals whose timescale is 1–2 orders of magnitude slower than the timescale of neural responses (tens–hundreds of milliseconds). Second, there is no integrated framework for mapping and quantifying spatiotemporal pRFs.

Here, we filled this gap in knowledge by developing a method for estimating spatiotemporal pRFs from fMRI. We combined a pRF mapping approach (Dumoulin and Wandell, 2008; Kay et al., 2013) with recent neural temporal encoding approaches (Stigliani et al., 2017, 2019; Zhou et al., 2018, 2019) to map and estimate spatiotemporal pRFs in each voxel in the visual system. To achieve the desired temporal estimates, we leveraged insights from recent fMRI studies that showed that not only stimulus duration, but also the number of transients and interstimulus intervals produce strong modulation of the amplitude of fMRI signals (Stigliani et al., 2017, 2019; Zhou et al., 2018, 2019). To measure spatiotemporal pRFs, we measured each voxel’s response to visual stimuli presented in different locations in the visual field under varying presentation timings (Fig 1). Then, we used a computational framework to estimate spatiotemporal pRF parameters in visual degrees and milliseconds from the fMRI response evoked by the stimulus (Fig 3). We estimated spatiotemporal pRFs in each voxel of multiple visual areas across three processing streams. The streams emerge in V1, continue to V2 and V3, and diverge into later visual areas in ventral (hV4, VO), lateral (LO, TO), and dorsal (V3AB/IPS) visual cortex.

**Figure 1.**
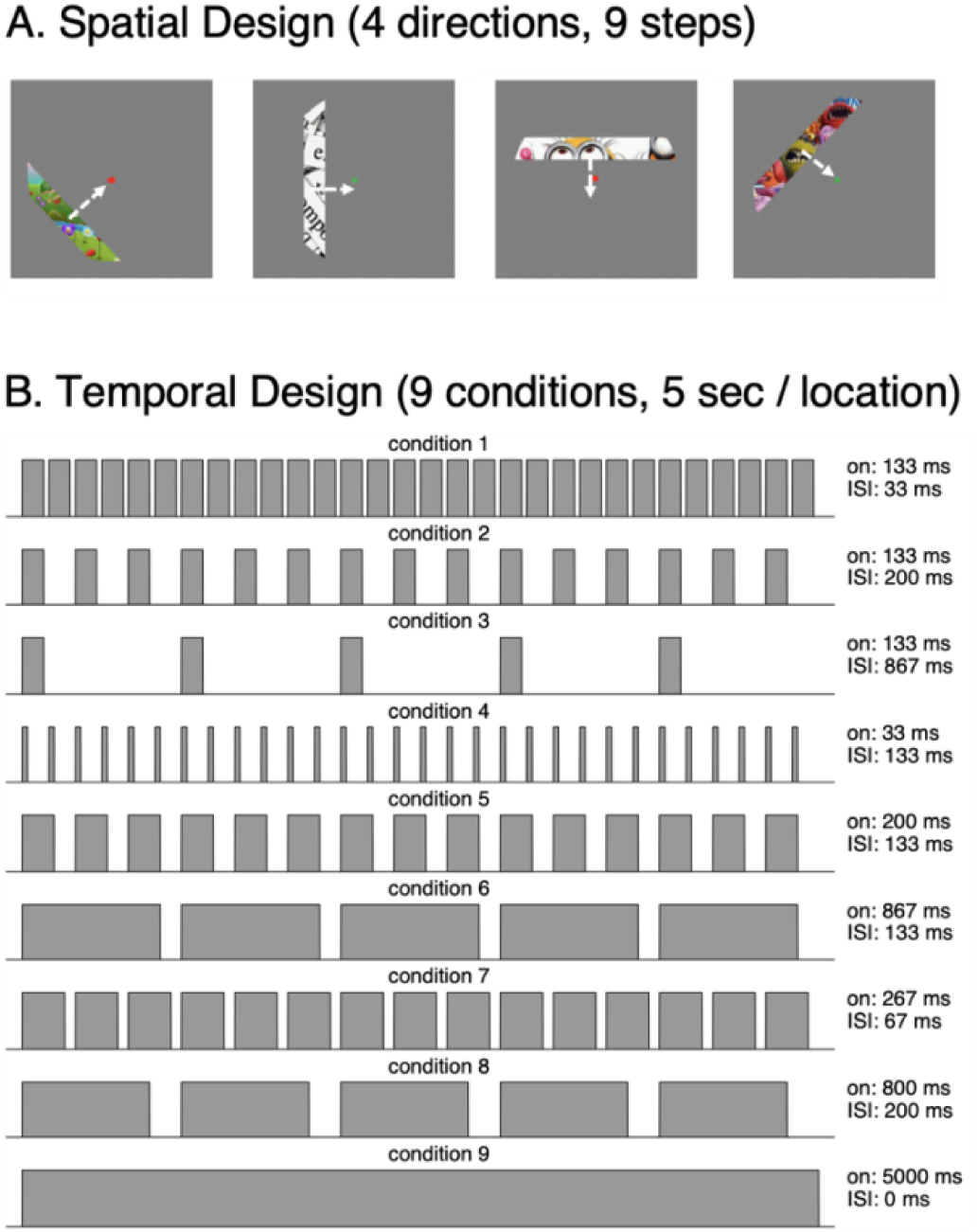
Spatiotemporal pRF experiment. In the experiment, participants viewed a flickering bar that swept the visual field while fixating and performing a color-change task at fixation. (A) Spatial design. A bar containing colorful stimuli continuously swept the visual field in 4 directions and each direction had 9 steps. Stimuli swept across a radius of 12° from fixation. Content of the bar was updated at a specific rate depending on temporal condition. (B) Temporal design. At each spatial location, the bar was presented for 5 seconds, in one of 9 different temporal conditions shown, where the bar content was updated with different random colorful cartoon snippets according to the temporal design of that condition. Each temporal condition was shown in each of the stimulus locations. *On*: on duration for each stimulus presentation. *ISI*: inter-stimulus interval. Example run: https://github.com/VPNL/stPRF#experiment

We examined how characteristics of spatiotemporal pRF may vary across visual areas. One possibility is that spatiotemporal pRFs vary across the visual processing hierarchy. Indeed, the spatial extent of pRFs progressively increases from early to later visual areas within a processing stream (Larsson and Heeger, 2006; Dumoulin and Wandell, 2008; Kay et al., 2013; Wandell and Winawer, 2015). Additionally, several studies suggest that temporal integration windows are larger in later than earlier visual areas (Hasson et al., 2008; Honey et al., 2012; Chaudhuri et al., 2015; Baldassano et al., 2017). These findings predict that pRFs in later visual areas will have both larger spatial and temporal integration windows (Zhou et al., 2018). Another possibility is that spatiotemporal pRFs vary across streams. Visual areas in the ventral stream (e.g., VO), that process static aspects of the stimulus may have pRFs with large spatial and large temporal integration windows (Van Essen and Gallant, 1994). In contrast, areas in the lateral stream (e.g., TO) that process motion information, may have spatiotemporal pRFs with large spatial but small temporal integration windows. These hypotheses are not mutually exclusive, as spatiotemporal pRFs may vary across both stages of the processing hierarchy and stream.

## Materials and Methods

### Participants

The study was approved by the Institutional Review Board of Stanford University. Prior to the start of the study, all participants gave written consent. Ten participants (ages 22 – 52 years, mean 30.6 years and s.d. 20.8 years; 7 female, 3 male). The demographics of participants were 4 East Asian, 3 white, 2 multicultural, and 1 Middle Eastern.

### Spatiotemporal pRF mapping Experiment

Participants performed 9 runs of the spatiotemporal pRF mapping experiment. While presented with the bar stimuli, participants were instructed to fixate on a central fixation point and perform a color-change detection task. Stimuli consisted of high contrast and colorful cartoon images (Finzi et al., 2021). To elicit a wide range of BOLD response profiles, we systemically varied the location and timing of stimulus presentation. Spatially (Fig 1A), each bar was created by dividing a cartoon image (radius of 12° visual angle) into 9 distinct apertures. Each bar had a width of 3° visual angle and there was a spatial overlap of 0.375° between adjacent bars. The 9 bars corresponded to the 9 steps in which the bar swept across the visual field in 4 different angles (0°, 45°, 90°, and 135°). Temporally (Fig 1B), the duration of each bar location was 5 s. Each 5-s bar location had one of the 9 different temporal conditions that varied in duration, inter-stimulus interval (ISI), and number of different stimuli. Specifically, temporal conditions 1, 2, and 3 had identical stimulus on durations of 133 ms per image with varying ISIs of 33 ms, 200 ms, and 867 ms, respectively. Temporal conditions 4, 5, and 6 had identical ISI durations of 133 ms with varying stimulus on durations of 33 ms, 200 ms, and 867 ms, respectively. Condition 1, 7, and 8 had an identical total stimulus on duration of 4 s while varying in the number images per bar location: 30, 15, and 5, respectively. Condition 9 had stimulus on duration of 5 s without an ISI, which served as a prolonged stimulus condition. The temporal conditions for each location were pseudo-randomly counterbalanced across runs and participants, making each run unique. Across the 9 runs, each temporal condition occurred once in each bar location. An example run of the experiment can be viewed online (https://github.com/VPNL/stPRF#experiment).

### Standard pRF mapping experiment

In a separate session, a traveling wave pRF mapping experiment was conducted to independently define borders of visual regions (Toonotopy) (Finzi et al., 2021). Specifically, we defined regions of interest (ROIs) which included: V1, V2, V3, hV4, VO (VO1 and VO2), LO (LO1 and LO2), TO (TO1 and TO2), V3AB and IPS (IPS0 and IPS1). This pRF mapping experiment used similar stimuli, the same visual field coverage (radius of 12° visual angle), number of angles (0°, 45°, 90°, and 135°) and task (color change detection task at fixation) as the spatiotemporal pRF mapping experiment. Different from the spatiotemporal pRF experiment: (i) images within each bar consisted of random cartoon images that changed at a constant rate of 8 Hz, (ii) bars were spatially less overlapping (0.27°), (iii) there were 12 steps in each direction, and (iv) each step had a 2-s duration. All participants completed 4 runs of the Toonotopy experiment.

### fMRI acquisition and preprocessing

During fMRI, stimuli were presented using an Eiki LC-WUL100L projector (resolution: 1920 x 1200; refresh rate: 60 Hz) using MATLAB (http://www.mathworks.com/) and Psychophysics Toolbox ((Brainard, 1997) http://psychtoolbox.org) MRI scanning was conducted on a 3T scanner (Signa, GE) with a Nova 16-channel head coil. Functional data were acquired using a T2*-weighted gradient-echo echo-planar imaging (GE-EPI) sequence (flip angle = 62°, TR = 1,000 ms, TE = 30 ms, field of view = 192 mm, 2.4 mm isotropic voxel size). Slices were prescribed to be parallel to calcarine sulcus to cover occipitotemporal cortex. A T1-weighted inplane image was collected for each participant, using the same prescription as the functional data, but higher resolution (0.75 x 0.75 x 2.4mm) to aid alignment to anatomical scan. A high-resolution anatomical scan (MPRAGE T1-weighted BRAVO pulse sequence, inversion time = 450 ms, flip angle = 12°, TE = 2.91 ms, 1 mm isotropic voxel size, field of view = 240 x 240 mm) were collected using a Nova 32-channel head coil. This anatomical scan was segmented in white and gray brain matter and used to reconstruct the cortical surface with FreeSurfer (version 6.0, (Fischl, 2012); http://freesurfer.net/).

All functional data were preprocessed using Vistasoft (http://github.com/vistalab/vistasoft) and SPM12 (https://github.com/spm/spm12). Functional images were aligned to each participant’s native space using T1-weighted inplane images. Then, the functional data were motion-corrected, and each voxel’s time courses were converted to percent signal change.

### Fitting hemodynamic response functions (HRF) for individual voxels

It has been well characterized that stimulus-evoked BOLD responses depend on both neuronal and hemodynamic properties (Lindquist and Wager, 2007; Polimeni and Lewis, 2021) and further, the hemodynamic responses may vary in response to different types of stimuli, among different regions of the visual cortex, and across individuals (Handwerker et al., 2004). All these factors may contribute to temporal parameter estimates in our model. In other words, if there are systematic variations in HRFs across different voxels and brain regions, using a single HRF for analysis may result in inaccurate estimates of spatiotemporal pRF parameters.

Thus, we performed an iterative linear fitting approach to estimate an optimized HRF for each voxel. First, we generated a stimulus design matrix for the Spatiotemporal pRF mapping experiment with 36 conditions (1 condition for each of the 9 bar locations and 4 orientations). Then, using a general linear model (GLM) approach, this design matrix was convolved with an HRF to generate predictors for each condition. For each iteration and for each voxel, the HRF parameters were optimized to minimize the difference between predicted fMRI time course and fMRI data. HRFs were parameterized as a sum of two-gamma functions (Friston et al., 1998) where each gamma function had two parameters: peak latency and full-width-half maximum (FWHM). The default Vistasoft HRF was also generated by using the two-gamma functions with peaks of 5.4 and 10.9 and FWHMs of 5.2, 7.35. Critically, the GLM design matrix coded the spatial location and orientation of the bar and disregarded the fine-grained temporal properties of the stimulus. Given that our experimental design included a wide range of temporal variabilities, the estimated HRFs are not biased to a specific temporal stimulus condition.

On average, the optimized HRFs across different ROIs were consistent across participants (Fig 2A, B). The estimated HRFs for all visual regions showed similar time to peak compared to the Vistasoft HRF (Fig 2A, dashed line), but some differences were observed including a delayed onset, wider width, and delayed undershoot. The across participant variability of estimated average HRFs were small (Fig 2A, B). However, when examining individual voxels’ HRFs within an ROI, we found a large degree of variation. As an example, while the average HRF profile for V1 and LO appeared similar (Fig 2B, C-black lines), there was substantial variability of HRF across voxels spanning these regions within a single participant (Fig 2C-colored lines). All results reported are with a voxel-wise optimized HRF unless otherwise stated.

**Figure 2.**
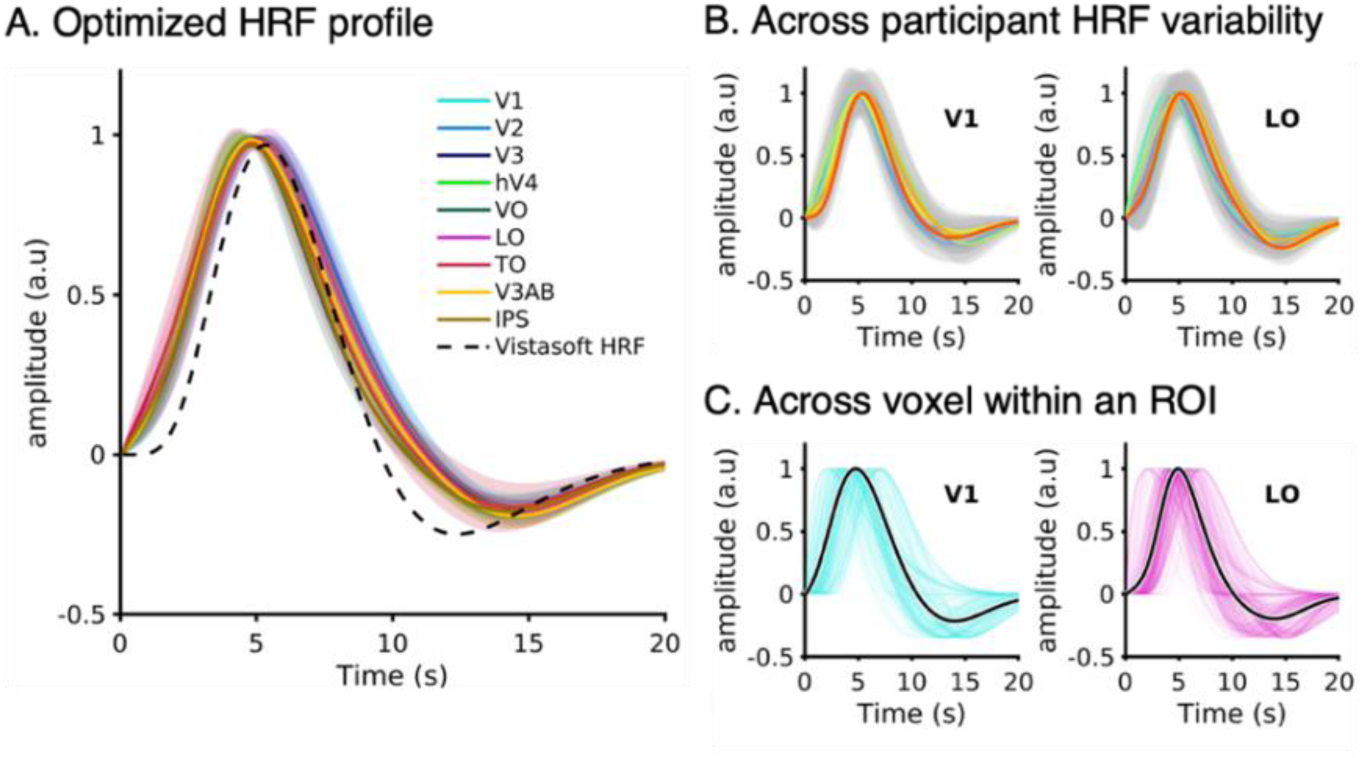
Optimized HRFs for individual voxels. For each voxel, we estimated its HRF using an optimization procedure. (A) The averaged estimated HRF across visual areas. The estimated HRFs for each visual area were averaged across voxels and participants. Heights of the HRFs are normalized to be 1 for visual comparison. *Blues:* V1, V2, and V3; *Greens:* hV4 and VO; *Reds:* LO and TO; *Yellows:* V3AB and IPS. *Shaded areas*: standard deviation across 10 participants. *Dashed line*: default HRF from Vistasoft. (B) To illustrate the across participant HRF variability, the mean HRF for V1 (left) and LO (right) is plotted for each individual participant. Each colored line indicates an individual participant. *Shaded gray area:* standard deviation across voxels for each participant. (C) To visualize within ROI variability of HRFs we show the HRFs of all voxels in V1 (left) and LO (right) from an example participant. Each line indicates the estimated HRF for a single voxel. *Solid black line:* the average HRF across voxels of that visual area.

### Spatiotemporal pRF modeling framework

#### Spatiotemporal framework (Fig 3A)

**Figure 3.**
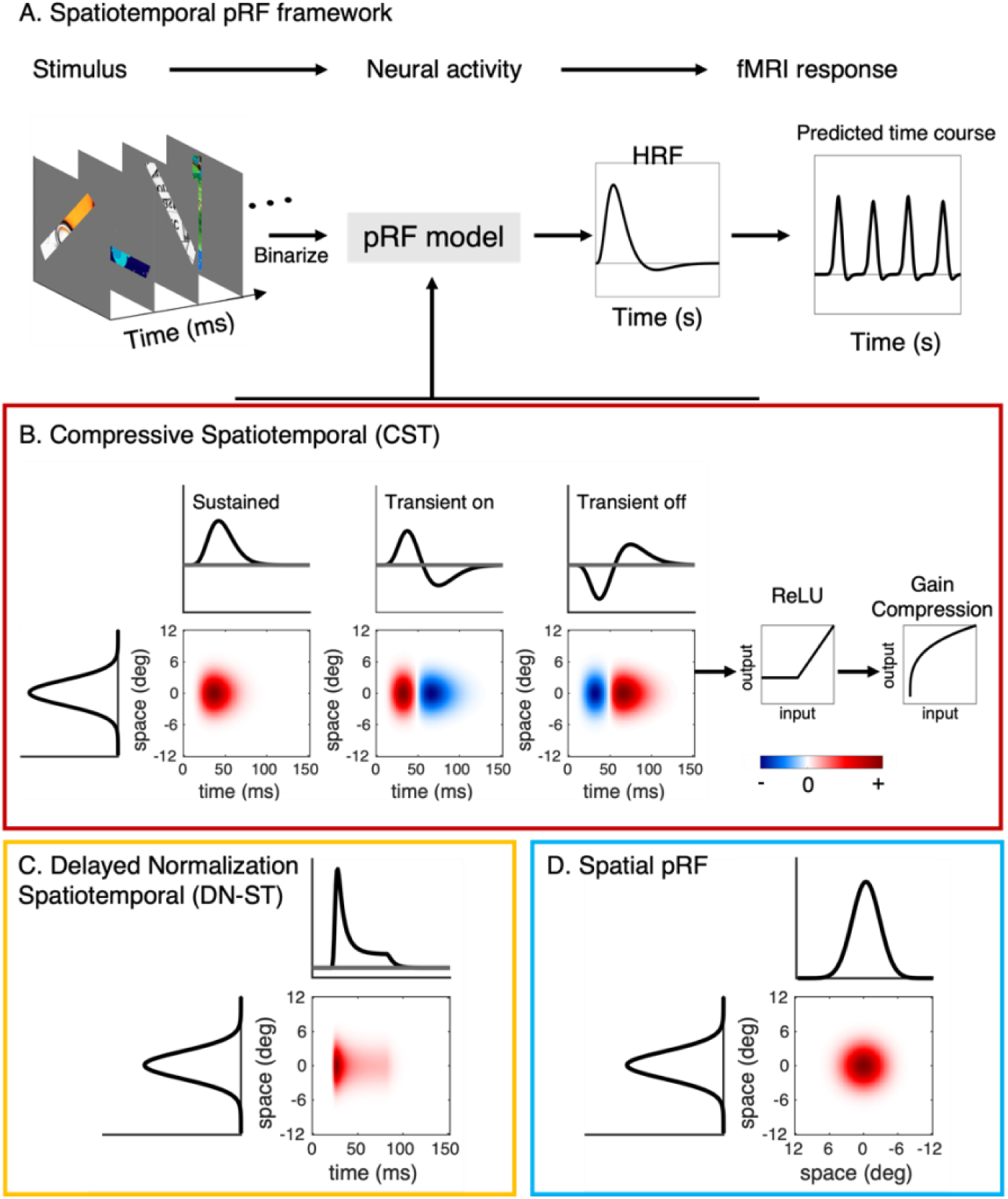
Modeling spatiotemporal population receptive fields. (A) Analysis pipeline of the spatiotemporal pRF framework. To predict the fMRI response in each voxel, the binarized visual stimulus is fed into a pRF model to predict the neural responses (temporal resolution of sequence and neural output is discretized into units of 10 ms) and then convolved with the hemodynamic impulse response function (HRF) and resampled to seconds. Unless otherwise stated, the HRF is the voxel-wise optimized HRF (Fig 2). We implemented and tested three pRF models: (B) Compressive spatiotemporal (CST) pRF model. The CST model consists of three spatiotemporal receptive fields that have an identical spatial receptive field (2D Gaussian) and three different temporal receptive field types: sustained (left), transient-on (middle), and transient-off (right). In each channel, the output undergoes rectification (ReLu) removing negative responses and compression by exponentiation. *Red:* positive signal amplitude (a.u.); *Blue:* negative signal amplitude (a.u.) For visualization, only the vertical spatial dimension is shown. (C) Delayed normalization spatiotemporal (DN-ST) pRF model. The spatial receptive field is a 2D Gaussian and the temporal receptive field uses a nonlinear impulse response function computed by rectification, exponentiation, and divisive normalization. (D) Spatial pRF model is a standard 2D Gaussian pRF; Here both spatial dimensions are shown.

The spatiotemporal pRF model is a stimulus-referred encoding model that predicts the BOLD response of each voxel while estimating both spatial and temporal neural characteristics of the pRF given a stimulus sequence. In general, an underlying assumption of the spatiotemporal pRF is that visual neurons integrate stimulus information over visual space and time (Adelson and Bergen, 1985; Watson and Ahumada, 1985).

First, we predicted neural activities from the stimulus and specific spatiotemporal receptive field model in centisecond resolution (Fig 3A). Depending on the model, this procedure involved linear and nonlinear computations. Then the predicted neural responses were convolved with a HRF and down sampled to 1 s to predict fMRI responses (Fig 3A). This step is linear and the same HRF was used across all models. Three pRF models were implemented: Compressive Spatiotemporal (CST, Fig 3B), Delayed Normalization Spatiotemporal (DN-ST, Fig 3C), and Spatial (Fig 3D).

This two-step stimulus to neural and neural to BOLD framework is theoretically and implementationally important. From the theoretical perspective, we sought to create a linking model that directly characterizes neuronal responses as well as their spatial and temporal nonlinearities and use it to predict the fMRI response to the stimulus. Implementationally, DN-ST and CST models apply non-linear operations at the neuronal stage while keeping a linear relationship between the predicted neural activity and BOLD response, as previous studies have shown that the temporal nonlinearities mostly arise from the neuronal activity to the stimulus. (Miller et al., 2001; Zhou et al., 2018).

#### Stimulus

The stimulus information is modeled in two spatial dimensions (*x*, *y*) and one temporal dimension (*t*) and referred to as *I*(*x*, *y*, *t*). Each frame of the stimulus sequence was binarized and resized to 61×61 pixels and the temporal resolution of stimuli sequences was 10 ms (centisecond). We implemented centisecond rather than a millisecond resolution to reduce computational time.

#### Spatiotemporal population receptive field (pRF) models

A spatiotemporal pRF is created by taking a pointwise multiplication of the neural spatial and temporal impulse response functions (Adelson and Bergen, 1985; Watson and Ahumada, 1985). To illustrate the spatiotemporal pRF profile in a 2D space, in Fig 3B-Cwe show a cross section of one spatial dimension (y-axis) with the temporal dimension (x-axis).

We used an identical spatial population receptive field (pRF) function for all three models, and only varied the temporal impulse functions specific to each model. The spatial pRF was modeled as a 2D isotropic Gaussian (Dumoulin and Wandell, 2008):

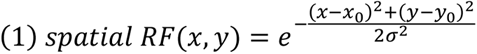

Where *x*_0_ and *y*_0_ are the center of the pRF in the visual field, and *σ* is the spatial extent of the pRF. All units are in degrees of visual angle (°). Fig 3D shows an example spatial pRF.

#### Compressive spatiotemporal (CST) pRF model (Fig 3B)

The CST model is inspired by neural measurements in macaque V1 (De Valois and Cottaris, 1998; De Valois et al., 2000; Conway and Livingstone, 2003) and human psychophysics (McKee and Taylor, 1984; Watson, 1986). The model consists of three spatiotemporal channels that have identical 2D Gaussian spatial receptive fields (spatial RF), each has a different neural temporal impulse response function: sustained, on-transient, and off-transient. The sustained impulse response function models the ongoing neural responses, while the transient impulse response functions computes changes in the neural response and highlights visual transitions. “On” and “Off” responses of the transient impulse response functions were separately modeled to account for increased neural responses with both stimulus onsets and offsets. These three spatiotemporal channels were designed to capture both prolonged and abrupt changes in neural responses to stimuli at a specific locations and sizes.

The CST model is implemented as follows: first, the dot product is applied between binarized spatiotemporal visual input *I*(*x*, *y*, *t*) and the *spatial RF* (equation 2). This procedure calculates the dot product between the stimulus and the spatial RF at each time point (effectively, the weighted sum of the spatial overlap between stimulus at time (*t*) and the pRF). Then, to predict spatiotemporal neural activity, we convolved the output with three different temporal impulse response functions *h_i_*(*t*), with *i* taking the values 1, 2, and 3. These temporal impulse response functions correspond to sustained *h*_1_(*t*), on-transient *h*_2_(*t*), and off-transient, *h*_3_(*t*) channels. Thus, the predicted spatiotemporal neural activity *r_i_*(*t*) for each of the channel can be expressed:

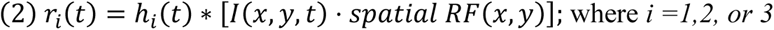

The sustained neural temporal impulse response was modeled by a gamma function:

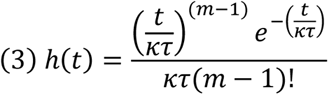

Where *t* is in msec, and *τ* is a fitted time constant. The values of *κ* and *m* parameters were same as previous studies (Watson, 1986; Stigliani et al., 2017, 2019), and were: *κ* = 1 and *m* = 9.

The on-transient impulse response function *h*_2_(t) was modeled as a difference between two gamma functions, which yielded a biphasic response. The excitatory component was the same as *h*_1_(t) and the inhibitory component was a gamma function with parameters: *κ* = 1.33 and *m* = 10. Note that, by definition, the peak of the sustained impulse response function is 1.33 times slower than the transient impulse response function. The transient-off impulse response function *h*_3_(t) is identical to *h*_2_(*t*), but with the opposite sign.

After spatiotemporal linear filtering, the CST model implements two nonlinearities in each channel: (i) a rectified linear unit (ReLU), and (ii) a compressive power-law exponentiation. The purpose of the ReLU is to rectify the negative component of the transient responses (the sustained response is always positive). Based on empirical observations. we reasoned that both on- and off-transient responses will increase neural firing rate and consequently increase BOLD signals. Nonlinear summation is thought to be a general sub-additive mechanism across spatial and temporal domains, as empirical data shows that the sum of responses to multiple stimuli over space is lower than the sum of responses to individual stimuli (Kay et al., 2013), and the response to prolonged stimulation is less than the response to multiple individual stimulation with the same overall duration (Zhou et al., 2018, 2019). The predicted neural activity of each channel (*i*) after nonlinear computations can be expressed as:

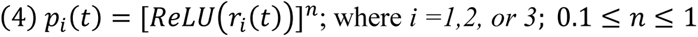

To predict fMRI responses resulting from sustained and transient neural responses, we convolved the predicted neural time courses *p_i_* (*t*) with the HRF and down sampled to 1 s resolution to match the temporal resolution of fMRI measurements. The predicted BOLD response is the weighted sum of the sustained and transient responses:

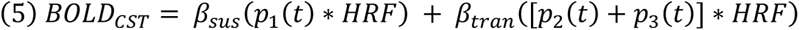

From fMRI data and using a general linear model (GLM) approach, sustained (*β_sus_*) and transient (*β_tran_*) scaling factors were estimated in each voxel (see also *Model fitting and parameter optimization*).

#### Delayed Normalization spatiotemporal pRF model (Fig 3C)

The spatiotemporal delayed normalization pRF model (DN-ST) has a 2D Gaussian spatial receptive field like the other models and a temporal impulse function that uses divisive normalization as well as an exponential decay function to model nonlinear neural temporal responses (Zhou et al., 2018, 2019; Groen et al., 2022).

Predicted neural activity of the DN-ST model is expressed as:

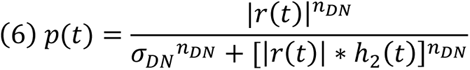

Where *σ_DN_* is a parameter for stimulus onset delay, and *n_DN_* is a scaling exponent. *r*(*t*) is the linear component of the neural response computed as the convolution between the neural temporal impulse response function *h*_1_(*t*) and the dot production of the stimulus *I*(*x*, *y*, *t*) with the *spatial pRF* (same as equation (2)).

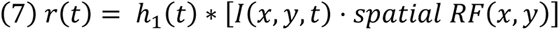

Following Zhou et al., 2019, the temporal impulse response function *h*_1_(*t*) was created using a gamma function with a time constant parameter *τ*_1_. We only implemented the monophasic version of the gamma function to reduce the number of free parameters.

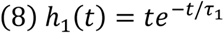

Another term in the denominator ℎ_2_(*t*) reflects the decay component and is implemented with a low-pass filter that uses an exponential decay function with a parameter *τ*_2_:

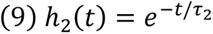

Finally, the predicted neural activity *p*(*t*) was convolved with the HRF to predict fMRI signals:

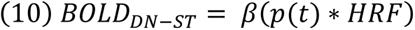

The *β*-weights approximate the magnitude of fMRI responses.

#### Spatial pRF model (Fig 3D)

The linear spatial pRF model is the same as (Dumoulin and Wandell, 2008). It implements the spatial pRF described in equation (1) to predict the neural response by computing the dot product between the stimulus and the spatial pRF. Then the neural response is convolved with the HRF to predict the BOLD response. The predicted BOLD response is:

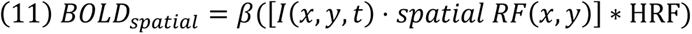

#### Model fitting and parameter optimization

The pRF parameters of each model at each voxel are determined by a two-stage coarse-to-fine approach that minimize differences between the predicted and measured fMRI time course. The first stage was a grid search procedure where we approximated the spatial RF parameters (x, y, and σ) for each voxel using the default Vistasoft HRF as in Dumoulin & Wandell (2008). The grid search procedure involved enumerating over combinations of potential spatial RF locations and sizes, where x and y were of a range that covered twice the size of the stimulus: 0° ≤ *x*, *y* ≤ 24°, 0.4° steps and *σ* was sampled up to the radius of the stimulus: 0.1° ≤ *σ* ≤ 12°, log-linear sampled, 96 steps. For the CST model, the grid search also included a range of compressive exponents: *n* = (0.25, 0.5, 0.75, and 1). Additionally, for the DN-ST and the CST models, we used a fixed set of default temporal parameters during the grid search: CST: *τ* = 4.93; DN-ST: *τ*_1_ = 0.05, *τ*_2_ = 0.1, *n_DN_* = 2, *σ_DN_* = 1 same as previous studies (Stigliani et al., 2019; Zhou et al., 2019)

In the second stage, a fine search was performed using a Bayesian Adaptive Direct Search (BADS) algorithm (Acerbi and Ma, 2017). To avoid local minima, we generated 3 sets of initial parameters using the estimated spatial parameters (x, y, and σ) from the grid search while randomly varying the other parameters. Here we fit all spatiotemporal parameters simultaneously. Lower and upper search bound of spatial parameters (x, y, and σ) were set to ±5° from the grid search estimation. The search range for remaining parameters were: DN-ST model: *τ*_1_: [0.01, 1], *τ*_2_: [0.1, 1], *σ*: [0.01, 0.5], *n*: [1, 6]; CST model *τ*: [4, 100], *n*: [0.1, 1]. Note that for the CST model, temporal (*τ*) and compression parameters (*n*) were identical across sustained and transient channels. For the fine search, different analyses used the default or optimized HRFs, and a GPU was utilized to accelerate computation time.

After performing the BADS optimization for each set, the parameter set that best explained the fMRI data was selected based on the highest variance explained (R^2^).

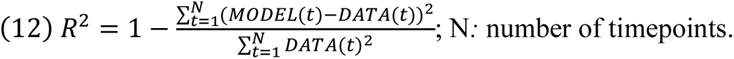

### Simulation software

We developed a simulation software to evaluate and ensure the computational validity of spatiotemporal pRF models. The simulation software was built to be compatible with previous validation software for fMRI BOLD responses (Lerma-Usabiaga et al., 2020). The simulation software has two components: (1) **Synthesizer:** generates synthetic fMRI time courses based on the stimulus and a specific pRF model. (2) **Solver:** Given a fMRI time course, a pRF model, and a stimulus sequence, the solver uses an optimization procedure to solve the pRF model parameters that best predict this time course.

To test the robustness of each of the spatiotemporal pRF models, we generated 300 different synthetic fMRI time courses for each model by randomly sampling a wide range of spatiotemporal pRF parameters. All three models shared identical ground-truth spatial parameters. Spatial receptive fields spatial parameters of *x* and *y* were randomly sampled from a normal distribution spanning an eccentricity range of 0°-10° and *σ* range of 0.2°-3° to match the stimulus aperture of our experiment. Additional parameters for the DN-ST and the CST model were randomly sampled from a uniform distribution with the same bounds used in the search algorithm described above. We used the default Vistasoft HRF for the simulation. After generating simulated time courses for each model and parameters, three types of additive noise (white noise, physiological, and low frequency drift noise) that are commonly found in fMRI signal were applied (Erhardt et al., 2012; Welvaert and Rosseel, 2014; Liu, 2016; Lerma-Usabiaga et al., 2020). The noise magnitude was systemically adjusted to match the typical range of signal-to-noise ratio (SNR) found in empirical fMRI data (SNR ≈ 0.1 dB; Fig 4B).

**Figure 4.**
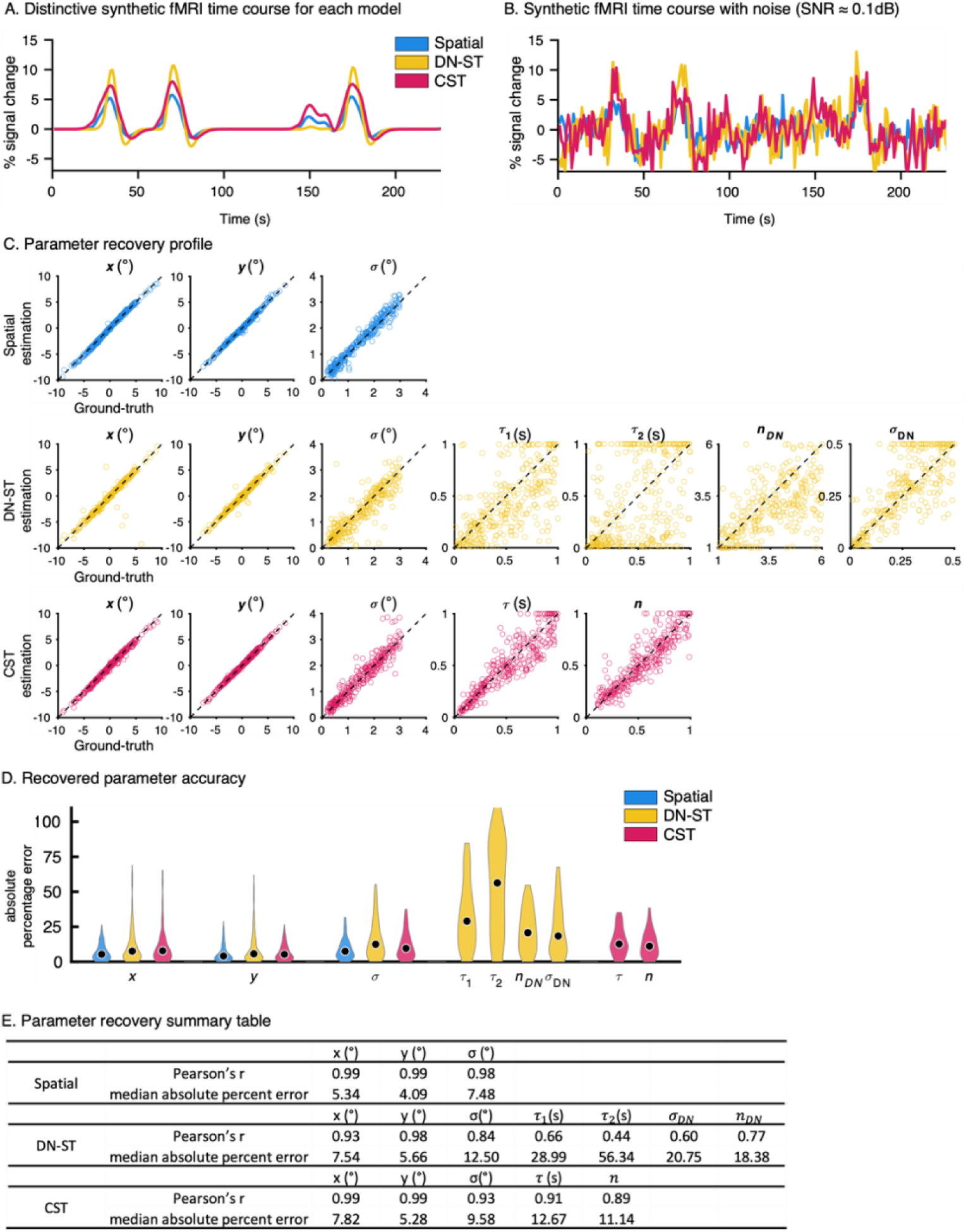
Simulator: Validating spatiotemporal pRF models. (A-B) Synthesizer. (A) An example time course of noiseless synthetic fMRI time courses generated with the same spatial parameters and the experimental paradigm of one run for three different pRF model types (blue: Spatial, yellow: DN-ST model, red: CST model). (B) Simulated time courses from (A) with added noise to simulate typical fMRI data. The amount of noise applied to each synthetic time course was matched across three models to yield mean SNR of 0.1 dB. (C-D) Solver Results. (C) Ground-truth parameter vs. estimated model parameters for 300 randomly generated pRFs for each model type. Each dot indicates a parameter recovery profile for one pRF from a noised added synthesized fMRI time course. Different model types have different numbers of parameters: CST model: 5 parameters (*x*, *y*, *σ*, *τ*, *n*). DN-ST model: 7 parameters (*x*, *y*, *σ*, *τ*_1_, *τ*_2_, *σ_DN_*, *n_DN_*), and Spatial model: three parameters (*x*, *y*, *σ*). The spatial parameters are in units of visual angle (°) and the temporal of *τ*, *τ*_1_, *τ*_2_ are in units of seconds (s). *Dashed line:* parameter estimate equals ground-truth. (D) Violin plots of the accuracy of each of the estimated parameters (absolute percentage error) for 300 simulated pRFs. *Black dots:* median value. For each parameter, outliers beyond 90^th^ percentile were excluded. (E) A table summarizing the accuracy of the parameters recovered compared to the ground-truth for 300 pRFs for each model. The median absolute percent errors are identical to the black dots in (D).

After generating synthetic fMRI time courses with noise, we tested whether each model can accurately recover its ground-truth spatiotemporal pRF parameters from the synthetized time courses. The parameters were solved using the two-staged coarse-to-fine approach as described above. The identical Vistasoft HRF that was used to generate the synthetic time courses was also used for solving. The performance of each spatiotemporal pRF model was evaluated by comparing the absolute percentage error between the solution and the ground truth parameters. The absolute percentage error was calculated by taking the absolute difference between the ground-truth and estimated parameter value, and then dividing by the ground-truth value. We also used Pearson’s (r) to compare parameter estimates of each model and the ground-truth parameter values.

### Estimating spatiotemporal pRFs across voxels of a region and across cortex

As different voxels’ pRF are centered in different locations in the visual field, to examine the aggregate properties of spatiotemporal receptive fields across voxels of a visual area, we zero-centered all pRFs spatial locations to x = 0 and y = 0. This allowed us to examine the distribution of spatiotemporal pRFs’ spatial and temporal windows in a region irrespective of their location in the visual field. These zero-centered spatiotemporal pRFs were then averaged across all voxels within each visual area in each participant and then across participants. For the CST model, this analysis was performed separately for the transient and sustained channels. Note that both sustained and transient impulse response functions share the same temporal parameter *τ*, and the time-to-peak ratio between the two impulse response functions are fixed. As the on- and off-transient spatiotemporal are identical except for their sign, for simplicity we only plotted the on-transient channel.

To assess the relationship between spatial and temporal windows of spatiotemporal pRF, we measured each voxel’s temporal window size by calculating the full-width-half-maximum (FWHM) of the sustained neural temporal impulse function. Then, we generated cortical maps of the pRF size and temporal window estimates. The map was generated for each participant and then transformed to FreeSurfer’s average cortical surface using cortex-based alignment and averaged across the 10 participants to generate the group average cortical map.

#### Statistical testing

To compare model performance, we used a 3-fold-cross-validation approach and averaged R^2^ across folds. In each fold, 2/3 of the data (6 runs concatenated) were used to estimate model parameters, and the left-out 1/3 of the data (3 runs concatenated) was used to estimate the variance explained by the model. The statistical significance of differences in model performance was computed by applying Fisher’s permutation test with 50,000 iterations and correcting the p-values for false discovery rate. To keep the same number of voxels for each model, if the analysis required comparison across different models, we did not exclude any of the fMRI voxels based on the variance explained. When examining the results of a particular model, we excluded voxels that account for less than 10% of the variance explained by that model. When reporting empirical fMRI results from the temporal parameter (*τ*), voxels with ill-posed estimates that are constrained at the boundary (*τ* = 100) were excluded.

Examining the relation between spatial and temporal windows: For each visual region, temporal window estimates were grouped into to 2-degree eccentricity bins. Then, linear mixed-effects analyses (LMM) with random participant intercept were conducted to explore the relationship between temporal window and eccentricity: temporalWindow ~ eccentricity + (1 | participant).

#### Data availability

The analysis code (https://github.com/VPNL/stPRF), data (https://osf.io/3gwhz/), and the pRF simulation software (https://github.com/vistalab/PRFmodel) are all accessible online.

## Results

### Validating spatiotemporal pRF models through simulation

To evaluate the robustness and validity of the spatiotemporal pRF modeling framework, we developed a simulation software with two purposes. First, to simulate experimental paradigms and test if they can be used to differentiate between predictions of different pRF models. Second, to validate the accuracy of the framework by testing if it recovers ground-truth pRF model parameters from simulated fMRI time courses with noise. By understanding how well our framework can recover parameters in simulated scenarios, we can define the scope of model interpretability given our experimental design and number of measurements.

### Synthesizer: different pRF models predict distinctive time courses to identical visual stimuli

Using the stimulus design in the spatiotemporal pRF mapping experiment (Fig 1), 300 noiseless and 300 noise-added synthetic fMRI time courses were generated for each of the pRF models: spatiotemporal pRF models (DN-ST and CST) and the conventional spatial pRF model (Dumoulin and Wandell, 2008). To compare how the temporal parameters affect fMRI time courses, synthetic fMRI time courses for each model using the same spatial parameters (*x*, *y*, *σ*) while varying model-specific parameters for DN-ST and CST.

We found that even with the same stimuli and identical spatial parameters, distinct synthetic fMRI time courses were generated for each model. An example of three noiseless synthetic time courses, one for each of the different pRF models, is shown in Fig 4A. All the example time courses had 4 distinct peaks at similar times during the experiment, corresponding to the 4 times the bar swept across the pRF. However, the different models generated distinctive fMRI time courses in terms of their amplitudes, latencies, and widths and these differences were preserved when typical fMRI noise was applied (Fig 4B). These different synthetic time courses confirmed that different types of pRF models can generate fMRI time courses that can be experimentally differentiated with our stimulus design.

### Solver: testing the accuracy and robustness of the spatiotemporal pRF framework vis-à-vis ground truth

We next tested whether the solver accurately recovers ground-truth pRF parameters from synthetic fMRI time courses. Specifically, using a synthetic fMRI time course with additive noise and a stimulus sequence as model inputs, the solver estimates the specific model parameters for each pRF. To evaluate the solver’s accuracy, we compared the estimated parameters to the ground-truth parameters that generated the synthetic time courses. Both the synthesizer and the solver used the same pRF model.

For noiseless synthetic fMRI time courses such as the ones in Fig 4A, the solver successfully recovered the spatiotemporal pRF parameters of all models with more than 99% accuracy. This result validates the analysis code and indicates that the solver can resolve pRF parameters from the planned experimental sequence for all model types.

When a typical level of empirical fMRI noise was added to the synthetic fMRI time courses, the models were still able to successfully recover the pRF parameters. The solver accurately recovered the spatial parameters (*x*, *y*, *σ*) (Fig 4C). Significant correlations were found between the predicted and ground-truth spatial parameters (Fig 4E, all correlations are significant p < .0001) across all the models. Estimated pRF center positions (*x*, *y*) were highly accurate (Fig 4C, estimated vs ground-truth along the dashed equality line). The median absolute percentage error (MAPE) for pRF center (x and y) was around 5.34% – 7.82% across the models (Fig 4D, 4E). The pRF size (*σ*) was also accurately recovered for all models (Fig 4C, 4D, 4E), yet with a higher MAPE of 7.48% – 12.5%. The higher accuracy of *x*, *y* than *σ* estimates is consistent with previous reports of lower accuracy in estimating pRF size than pRF center positions (Lage-Castellanos et al., 2020; Lerma-Usabiaga et al., 2020).

Although the solver was overall less accurate for estimating the temporal parameters than spatial parameters from noisy synthetic data, it successfully recovered the temporal parameters for the DN-ST model (*τ*_1_, *τ*_2_, *σ_DN_*, *n_DN_*) and the temporal (*τ*) and compression (*n*) parameters from the CST model, with better estimates for the CST than the DN-ST model (Fig 4C, 4D; all correlations between estimated and ground-truth parameters are significant p < .0001). Remarkably, the CST model was able to resolve the millisecond temporal receptive field parameter (*τ*) with a MAPE < 13% (Fig 4D, 4E). Despite the sluggish nature of hemodynamic responses, the stimulated result suggests that given a ~200 ms temporal receptive field, we would only find a ±26 ms margin of error. The compression parameter, *n*, was also reliably estimated. The estimates of the DN-ST model temporal parameters were also significantly correlated with the ground-truth temporal parameters (Fig 4C, 4E, p < .0001), but the estimates were less accurate than the CST model, with *τ*_2_ being the least accurate parameter (Fig 4D, Fig 1E). This variability of *τ*_2_ estimates may be due to an insufficient number of conditions with prolonged presentation, as the *τ*_2_ parameter mainly controls the temporal decay rate of the response to prolonged stimulus durations.

### Spatiotemporal pRF models outperforms spatial only pRF model

We next assessed how well the three pRF models: Spatial, DN-ST, and CST, predicted the empirical fMRI time courses in the spatiotemporal mapping experiment (Fig 1) in each of our 10 participants. Note that the time courses predicted from the spatial model depend only on the location and duration of the stimulus, whereas the predictions of the DN-ST and CST models also depend on the spatiotemporal aspects of the stimulus at each location, resulting in more complex fMRI responses.

The spatiotemporal stimulus design evoked a wide range of BOLD time courses in voxels of the visual system. Based on each voxel’s fMRI time course, pRF parameters for each of the three models were estimated. Results show that fMRI responses at the voxel level were well captured by the DN-ST (example voxel, Fig 5A-yellow) and CST (example voxel, Fig 5A-red) models. However, the spatial model failed to predict the fMRI time courses in multiple ways. For example, as shown in the example V1 voxel in Fig 5A-blue, the Spatial model substantially under predicted responses for 4^th^ and 5^th^ peaks, over predicted responses for the 1^st^ and 2^nd^ peak, and failed to predict the widths of the 3^rd^ and 7^th^ peaks. The failures of the Spatial pRF model mostly occurred in stimulus conditions with fast temporal rates. In contrast, the DN-ST and CST models were able to predict both the amplitude and width of these peaks that the Spatial model failed to predict.

**Figure 5.**
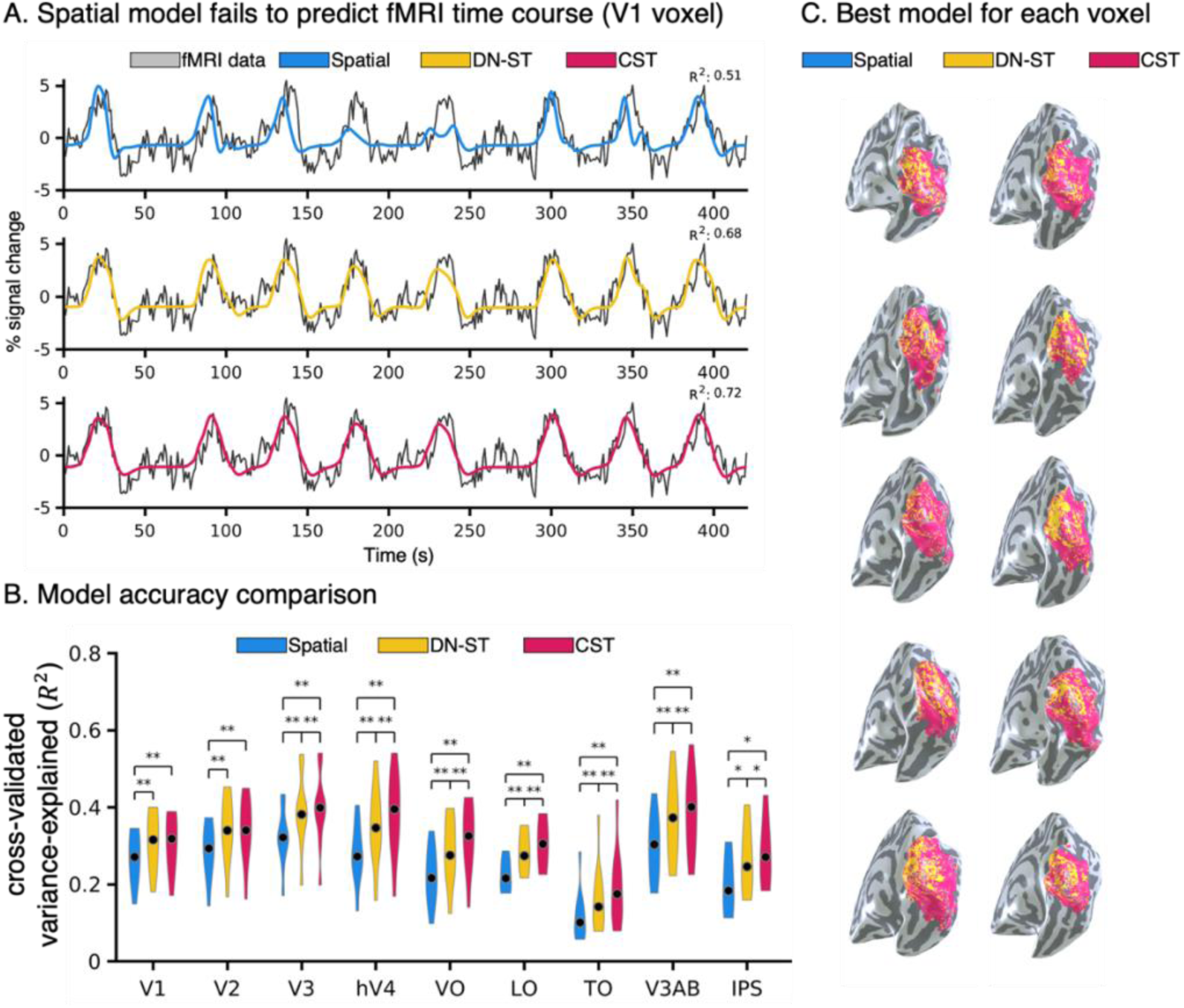
fMRI data: comparing spatiotemporal pRF models to fMRI voxel data. (A) *Black:* An example fMRI time course from an example V1 voxel. *Colored lines:* model predictions; *blue*: Spatial, *yellow:* DN-ST, *red:* CST. The same fMRI data is shown in all rows. (B) Model accuracy comparison. Violin plots of average cross-validated variance explained (R^2^) across each participant’s voxels and area for each of three model in nine retinotopic visual areas spanning the ventral, lateral, and dorsal processing streams. *Black dots:* median value. *Asterisk:* Significantly different than spatial model, permutation test, * p < .05, ** p < .01. (C) The best performing model (R^2^) for each voxel. Data are shown for each of the ten participants on their inflated right hemisphere. *Blue:* best model is Spatial, *yellow:* best model is DN-ST, *red*: best model is CST.

We quantified how well the various models predicted the experimental time courses in each voxel using a 3-fold cross-validation approach. On average, the Spatial, DN-ST, and CST models accounted for 25%, 30%, and 33% of the variance, respectively. Each model’s accuracy across multiple retinotopic visual areas is plotted in Fig 5B. The DN-ST and CST model outperformed the Spatial model in predicting fMRI responses in single voxels of all tested visual areas. Indeed, the 3-fold cross-validated variance explained (R^2^) by the DN-ST and CST models was significantly higher than the spatial model (Fig 5B; permutation test, V1, V2, V3, hV4, VO, LO, TO, V3AB, p < .01; IPS, p< .05). Comparing the spatiotemporal pRF models, the CST model outperformed DN-ST model across all visual regions (permutation test, V3, hV4, VO, LO, TO, V3AB, p < .01; IPS, p < .05) except V1 and V2, where performance did not differ significantly between the two spatiotemporal models.

To examine if there were systematic and spatial differences in how well models fit the data across cortex, we numerically compared the variance explained by the three pRF models in each voxel. Then, we visualized on the cortical surface which model best predicted the response of each voxel (Fig 5C). We found that: (i) in all participants, there were almost no voxels for which the Spatial model was the best model (very small number of blue voxels in Fig 5C), (ii) for most of the voxels the CST model was the best (except for voxels around the occipital pole for which the DN-ST model was better), and (iii) consistent with the qualitative analyses, the advantage of the CST vs. DN-ST models was more pronounced in later visual areas.

Together, this analysis: (i) supports our hypothesis that both spatial and temporal aspects of the stimulus contribute to fMRI signals at the voxel level, (ii) shows that both DN-ST and CST spatiotemporal pRF models can capture spatial and temporal dynamics of neural responses, and (iii) suggests that considering the nonlinear temporal dynamics, both at the stimulus and neuronal level in millisecond resolution into a pRF model is crucial for accurately predicting fMRI timeseries at the voxel level.

### Spatial estimates are consistent across models

As spatiotemporal pRF models are newly developed, it is important to test if the spatial pRF parameters they estimate are topographically organized and replicate well-established retinotopic maps in the human visual cortex. Thus, we next used the estimated spatial parameters to generate phase, eccentricity, and pRF size maps for each participant and model (Fig 6A, B, C). We also quantitatively compared the estimated pRF position (polar angle), eccentricity (°), and size (*σ*) for the DN-ST and CST pRF models to those of the Spatial model (Fig 6D, E, F).

**Figure 6.**
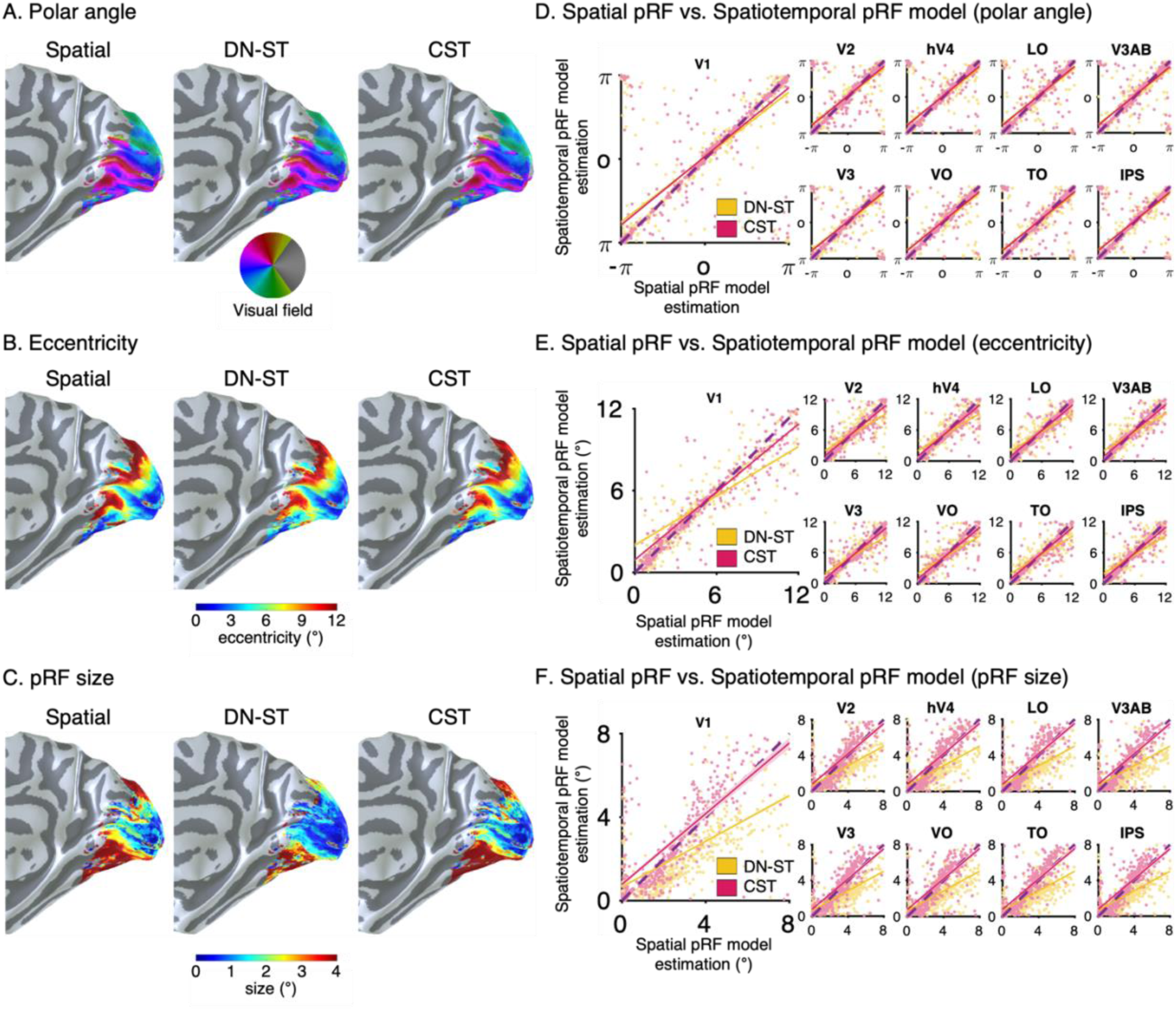
Spatial pRF parameter estimated for the spatial, DN-ST, and CST models are similar. (A) Polar angle, (B) eccentricity, and (C) pRF size maps on the right inflated cortical surface of an example participant for each of the three models. (D-F) Scatter plot comparing pRF from spatiotemporal pRF models (*yellow:* DN-ST and *red:* CST, y-axis) to estimates from spatial pRF model (x-axis) for each of nine visual aeras. *Colored yellow and red lines:* linear regression fits. Each dot indicates the parameter estimate for one voxel. For visualization, we randomly sampled 50 voxels from each participant for each ROI. *Dotted black line:* reference line of identical estimates. (D) Polar angle. (D) Eccentricity. (F) pRF size (*σ*).

We found that all three models produced robust and consistent estimates of each voxel’s pRF positions. From pRF center positions (*x*, *y*) we generated phase and eccentricity maps, which were indistinguishable across the three pRF models and had the expected phase reversal topography (see example participant, Fig 6A, B). Moreover, all models generated similar pRF size estimates with the expected topography with increasing sizes along posterior-anterior axis (Fig 6C). Although in high agreement, pRF size estimates were more variable across models (Fig 6F). In all tested visual areas, the Spatial model and CST model produced similar pRF size estimates (Fig 6F, red), while the DN-ST model consistently estimated a smaller spatial pRF size compared to that estimated by the spatial model (Fig 6F, yellow). The reliable estimate of spatial pRF properties across models suggests that the incorporation of additional temporal parameters into the pRF model does not reduce the statistical power to map spatial pRFs in the human visual cortex.

### Differences in temporal estimates cannot be explained by differences in HRFs

We next characterized the temporal parameters of spatiotemporal pRFs across voxels of the visual system. Under our experimental paradigm, the simulations show that temporal parameters of the CST model were more accurately recovered than the DN-ST model (Fig 4C, D) and experimental data revealed that the cross-validated variance explained of fMRI responses was higher for the CST than DN-ST model for most voxels (Fig 5C). Thus, in the following sections we report the temporal and spatiotemporal characteristics estimated from the CST model. In the CST model, the temporal parameter *τ* governs time-to-peak of the neural temporal impulse response function, which approximates both the response latency and temporal integration window of neural responses.

We estimated the neural time-to-peak (τ) in each voxel and visual area, then plotted the distribution of neural time-to-peak across voxels of an area averaged across subjects for early visual areas (V1, V2, V3) as well as intermediate and later visual areas in the ventral (V4, VO), lateral (LO, TO), and dorsal streams (V3AB, IPS). As evident from Fig 7A, neural time-to-peak increases from earlier to later areas but also varies within each visual area.

**Figure 7.**
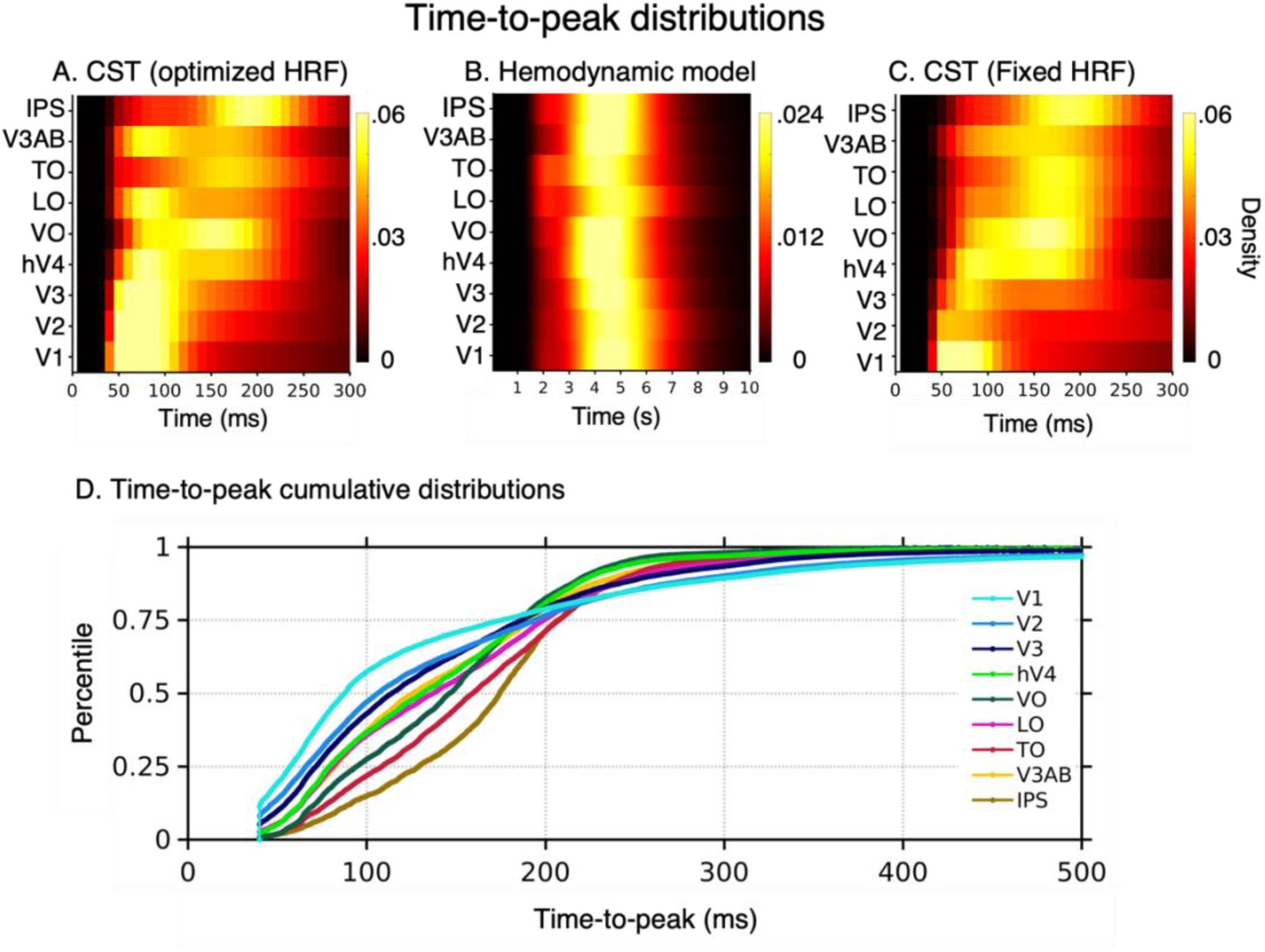
Temporal pRF estimates from the CST model. (A-C) Each row shows a voxel-wise time-to-peak distribution for each visual area. The distributions were averaged across participants within each ROI. (A) Time-to-peak distribution estimated from the CST model, using an optimized HRF for each voxel. (B) The distribution of optimized HRF time-to-peak. Note that unlike the CST model, the temporal resolution for the hemodynamic model is in seconds. (C) Time-to-peak estimated from the CST model using a single default Vistasoft HRF for all voxels. (D) The cumulative time-to-peak distribution across visual areas from the CST model with optimized voxel-wise HRF for each of nine visual areas. The datapoints from all participants were concatenated for each visual area.

As we estimated for each voxel its optimized HRF (Fig 2), it is possible that this variability in the time-to-peak is of hemodynamic rather than neural origin. To examine this possibility, we conducted two additional analyses. First, we analyzed the distribution of time-to-peak of the hemodynamic functions within and across areas. We reasoned that if the temporal variability is of hemodynamic origin, then the distributions of optimized HRF time-to-peak will mirror the distributions of the estimated neural time-to-peak. However, this was not the case. The distribution of hemodynamic time-to-peak ranged from 3-6 seconds and was similar across all tested visual areas Critically, this distribution did not show the between-area differences in neural time-to-peak estimated by the CST model (Fig 7B). Second, we repeated the calculations of the CST neural time-to-peak (τ) using a fixed HRF for all voxels (Vistasoft default HRF). We hypothesized that if the estimated neural time-to-peak interacts with the estimated HRF parameters for each voxel, then using a fixed HRF will qualitatively change the estimates of neural time-to-peak within and across areas. However, we found that using a fixed HRF for all voxels produces quantitative but not qualitative changes in estimates of neural time-to-peak (Fig 7C). Specifically, using a fixed HRF increased the within-area variability of neural time-to-peak estimates (e.g., compare V3 in 7C to 7A), but did not change the observation that the time-to-peak progressively increases across visual the hierarchy. These analyses give us confidence that our approach enables estimating neural temporal properties. In the next sections, we will examine in detail the temporal and spatiotemporal parameters from the CST model with the voxel-wise optimized HRF.

### Hierarchical temporal latency delay across visual streams

An open question is how the time-to-peak and processing time window changes across the visual hierarchy and processing streams. A feedforward model of the visual system predicts that the time-to-peak will progressively increase across the visual hierarchy (Nowak and Bullier, 1997). Additionally, the ventral stream which is associated with perception of invariant properties of objects and people (such as their identify or category) may be slower and have longer times-to-peak than other streams, like the lateral stream, which is thought to process motion and dynamical aspects of the visual scene (Van Essen and Gallant, 1994; Weiner and Grill-Spector, 2013; Pitcher and Ungerleider, 2021; Wurm and Caramazza, 2022).

We find differences in time-to-peak both across the hierarchy and across streams, thus finding evidence for both hypotheses. Across the hierarchy, time-to-peak increases from earlier (V1, V2, V3) to later visual areas in each processing stream generating a cascade of neuronal signal propagation. We find that voxels in V1 had the earliest time-to-peak latencies (*τ* ~ 50 − 110 ms, Fig 7A), and that time-to-peak was less than 100 ms for more than 50% of V1 voxels (Fig 7D). V2 followed V1 (*τ* ~ 50 − 120 *ms*), and V3 followed V2 (*τ* ~ 50 − 130 *ms*). In the ventral visual stream, later time-to-peak was observed in voxels of intermediate visual area hV4 (*τ* ~ 70 − 170 *ms*, Fig 7A, D-green) which were followed by responses in a late visual area, VO (*τ* ~ 80 − 200 *ms*, Fig 7A, D-dark green). Similar trends were observed in the lateral (LO to TO, pink and red in Fig 7D) and the dorsal streams (V3AB to IPS, yellow and brown in Fig 7D). Timing of the intermediate areas did not vary across streams with hV4 (ventral), LO (lateral), and V3AB (dorsal) having similar time-to-peaks. However, we find differences across later areas of the different streams. TO (lateral) had longer times to peak than VO (ventral), and IPS had the latest time-to-peak (*τ* ~ 160 − 230 *ms*) compared to other tested visual regions (Fig 7A, D-brown).

### Spatiotemporal processing across visual streams

Crucially, the CST model allows us to examine the spatiotemporal properties of pRFs across visual streams. Prior research has suggested that both spatial receptive field sizes (see review, (Wandell and Winawer, 2015) and temporal windows (Hasson et al., 2008; Honey et al., 2012; Murray et al., 2014; Baldassano et al., 2017) progressively increase across the visual hierarchy. However, before the current framework it was not possible to estimate spatial and temporal population receptive field sizes in degrees and milliseconds directly at the voxel level.

Fig 8A shows the average sustained (top) and transient (bottom) spatiotemporal pRF across voxels of a visual area. Earlier visual areas had smaller pRF sizes and shorter temporal windows compared to later visual areas for both sustained and transient channels. For example, ascending the ventral stream (V1->V2->V3->hV4->VO), spatiotemporal pRF integration windows progressively increased both spatially and temporally (Fig 8B). In the sustained channel, V1 pRFs integrate visual information spatially across 0.41±0.05° (median±SEM) and temporally across 58.77±10.71ms, hV4 across 1.63±0.13° and 88.27±9.93ms, and VO across 2.39±0.18° and 98.19±9.66ms (Fig 8A, B). Likewise, in the lateral stream, TO spatiotemporal pRFs were larger in space and time compared to LO, and in the dorsal stream, IPS spatiotemporal pRFs were larger in space and time than V3AB. In the transient channel, we find a similar progression (Fig 8A, bottom).

**Figure 8.**
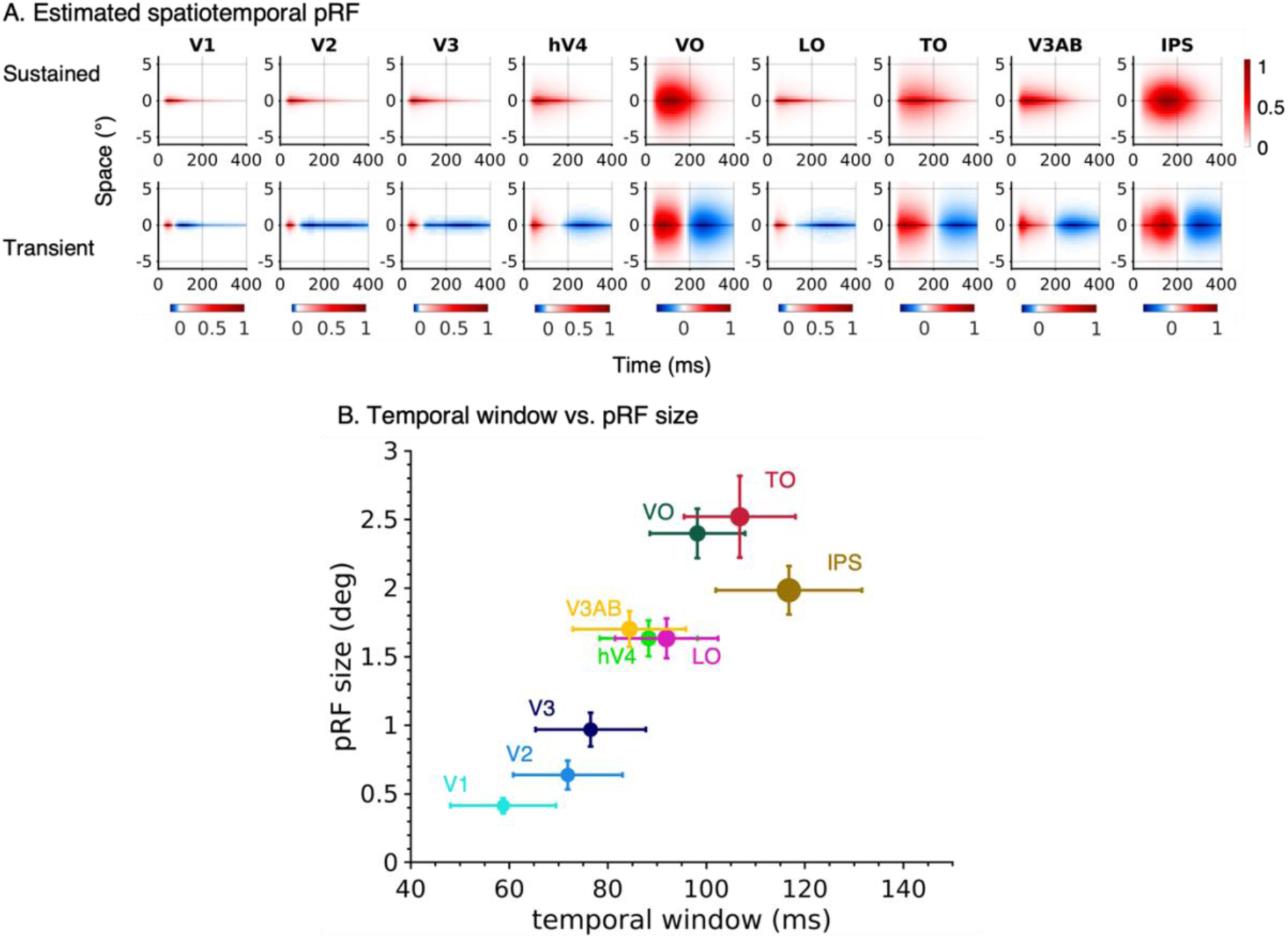
Spatiotemporal pRF characteristics across nine visual areas spanning three visual streams. (A) Spatiotemporal pRF profiles for sustained and transient (on) pRFs were estimated for each voxel and averaged across participants for each visual area. The spatial location of all pRF was zero-centered to (0,0) before averaging. The x-axis represents time (ms), and the y-axis represents a cross-section of visual space (°). The red-white-blue colormap is the average of spatiotemporal pRFs. (B) pRF temporal window (FWHM of the sustained temporal impulse response function) versus pRF size (*σ*, deg). *Colored dots:* median values for each visual area. Size of colored dots is related to median values of spatiotemporal compression exponent *(1/n*). *Error bars:* ±1SEM across participants in each dimension.

To quantitively compare the spatiotemporal pRF size and temporal window estimates, we computed the average pRF size and average temporal window of each visual area (y-dimension of Fig 8A, top). The comparison showed a positive relationship, revealing a general trend of progressively increasing spatial and temporal windows along the visual hierarchy. Specifically, spatiotemporal pRF size and temporal window progressively increased from V1 to V2 to V3 (Fig 8B-blues). Intermediate areas of all three visual streams (hV4, LO, and V3AB) had similar spatial and temporal pRF sizes which were larger spatially, and longer temporally than V1-V3. The spatiotemporal properties of later visual areas (VO, TO, and IPS) differed across streams. Notably, spatiotemporal pRFs in VO, a ventral area, were smaller spatially and shorter temporally (Fig 8B-dark green) than pRFs in TO, a lateral area (Fig 8B-red), whereas IPS had pRFs that were smaller spatially, but longest temporally (Fig 8B-brown). In addition, the amount of spatiotemporal compression increased along the visual hierarchy (Fig 8B, marker size ~ 1/*n*). The compressive exponent (*n*) in V1: 0.28±0.03, V2: 0.25±0.06, V3: 0.24±0.06, hV4: 0.22±0.04, VO: 0.21±0.03, LO: 0.19±0.02, TO: 0.18±0.02, V3AB: 0.21±0.02, IPS 0.14±0.02, which is comparable to the increase in spatial compression levels from early to late visual areas (Kay et al., 2013). These results illustrate an interesting coupling between spatial and temporal receptive field integration windows that progressively increase within each of the three processing streams and diverge across streams in the later visual areas.

Finally, we investigated the spatial topography of spatiotemporal receptive fields across visual cortex. Qualitatively examining the maps of temporal window and pRF size reveals a topographic coupling of spatial and temporal pRF windows. We first examined this topography in early visual areas (Fig 9A-left), where we observed that pRF temporal integration windows (Fig 9A-top-left) increase along a posterior to anterior gradient and pRF sizes (Fig 9A-bottom-left) also increase along the same axis. Indeed, a linear mixed model (LMM) revealed a significant relationship between temporal window and eccentricity in V1, V2, V3 (Fig 9B), whereby time integration windows were progressively larger in more peripheral eccentricities (V1 (t(58) = 7.66, p < .001), V2 (t(58) = 6.25, P < .001), and V3 (t(58) = 4.09, P < .001 with corresponding beta coefficients of 0.4, 0.35, and 0.27, respectively). Despite some individual variability in the intercept among participants, this relationship is visible in almost all participants. As eccentricity also increases from posterior to anterior in early visual areas, this suggest that in V1-V3, more eccentric pRFs have larger spatial and temporal integration windows than foveal pRFs. Fig 9C visualizes the average spatiotemporal pRFs at two eccentricity bands: 0-4° and 4-12°. Indeed, in V1-V3, both pRF size and temporal window were larger in eccentricities exceeding 4° than the central 4°. The proportional increases in both spatial and temporal dimensions with eccentricity suggests that there are computational similarities in how the visual system encodes spatial and temporal information at the center and periphery.

**Figure 9.**
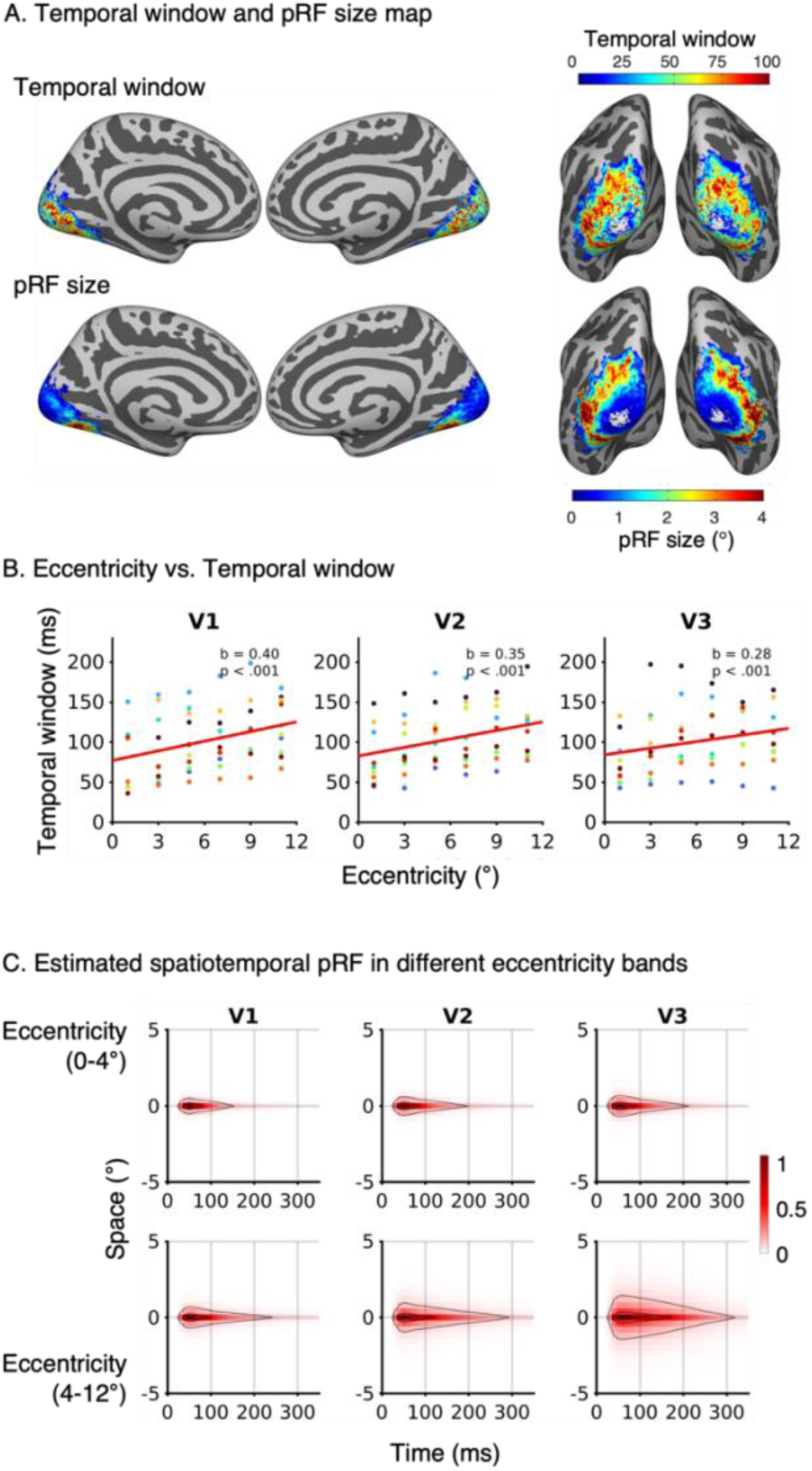
(A) Average pRF temporal window and spatial window on the FreeSurfer average brain. Both hemispheres and two different viewpoints are shown (*left:* medial view, *right:* posterior view). *Top:* FWHM of average sustained temporal window estimates; *Bottom:* average pRF size. (B) The relationship between eccentricity and temporal window in V1, V2, and V3. *Colored dots*: individual participant. *Solid line:* fitted LMM regression line. (C) Mean sustained spatiotemporal pRF in two different eccentricity bands (0-4° and 4-12°). Contour lines indicate 10^th^, 50^th^, and 90^th^ percentile levels.

A similar topography showing a coupling between pRF size and temporal window is also seen in the lateral surface where more posterior regions have smaller spatial and temporal integration windows than anterior regions (Fig 9A-right). Within higher-level areas eccentricity-dependent relation between pRF size and temporal windows were significant in LO (t(58) = 2.31, p < .05), while the opposite effect was observed in VO (t(58) = −0.46, p < .001). No significant eccentricity-dependent effects were found in V4 (t(58) = −0.48, p = 0.63), TO (t(58) = −0.63, p = 0.54), V3AB (t(58) = 1.95, p = 0.056), and IPS (t(58) = −1.53, p = 0.13).

Together these analyses reveal two organizational principles. First, both spatial and temporal integration windows progressive increase across the visual hierarchy. Second, within early visual cortex, both spatial and temporal integration windows increase with eccentricity.

## Discussion

Here we developed a new mapping and computational framework to estimate spatiotemporal pRFs in each voxel using fMRI. We find that spatiotemporal pRFs with a Gaussian spatial profile, which includes sustained and transient temporal channels, and a compressive nonlinearity better explain fMRI responses than conventional Gaussian pRFs. This is because the conventional Gaussian pRFs lack explicit temporal functions and instead linearly sum over stimulus location and duration. Spatiotemporal pRFs’ spatial parameters (pRF center and size), and their cortical topography replicate findings from conventional pRF methods (Wandell and Winawer, 2015). Temporal parameters (peak-latency and temporal integration windows) and compressive nonlinearities progressively increase from earlier to later areas of visual streams. Interestingly, we find spatiotemporal coupling whereby pRFs in intermediate and later visual areas have both larger spatial and temporal integration windows than pRFs in earlier visual areas. Together, this spatiotemporal pRF framework pushes the temporal limits of fMRI to understand how spatiotemporal information is encoded in visual degrees and milliseconds across neural populations in human visual cortex.

### Spatiotemporal pRF Framework

The spatiotemporal framework we developed serves as a testbed for systematically comparing pRF models. The standardized format of the spatiotemporal framework ensures model reproducibility and validity, as all the models are tested and compared using matched HRF, optimization algorithm, and SNR of synthetic time courses. Moreover, each step of the data analysis pipeline is modularized to facilitate the examination of individual components.

Through simulation, we defined the scope of model parameter interpretability, given a unique visual input, number of measurements, and expected level of fMRI noise. This quantification enables us to validate that the solved model parameters accurately recover the underlying spatiotemporal pRF from the stimulus and synthesized fMRI response. Applying the spatiotemporal pRF approach to both synthesized and measured fMRI data revealed that spatial parameter estimates were similar across the three pRF models tested (Spatial, DN-ST, and CST), and temporal parameter estimates of the DN-ST model were noisier than the CST model. Future research can leverage the simulator to design optimal mapping sequences for different spatiotemporal pRF models, which would provide opportunities for better estimating model parameters (e.g., temporal parameters of DN-ST) and comparison between models. Additionally, the spatiotemporal pRF model can be expanded to encode additional aspects of the stimulus such as contrast, shape, and motion to models additional aspects of cortical processing that vary across space and time.

### Differences in HRF temporal parameters cannot explain neural temporal dynamics across visual areas

The spatiotemporal pRFs approach first predicts neuronal responses to the stimulus in milliseconds and then convolves the predicted neural response with the HRF to predict BOLD signals. This implementation allowed us to separately evaluate the effect of neural and hemodynamic responses on the resulting BOLD signal. Our data revealed that neural temporal parameters varied systematically across visual areas and optimized HRF parameters varied more across voxels within an area than across areas (Fig 7B). Critically, this variability in HRF parameters did not explain our neural estimates as neural temporal parameters across areas were similar across models solved with optimized HRFs and constant HRF functions (Fig 7A, C). Nonetheless, the optimized HRF reduced between-voxel variability consistent with (Prince et al., 2022). Notably, this spatiotemporal pRF approach is not only necessary for predicting the observed BOLD responses – especially to brief stimuli that are separate by short interstimulus intervals – but also is a more parsimonious approach than a schema of different HRFs for fast and slow stimuli (Lewis et al., 2016; Polimeni and Lewis, 2021).

### Comparing human temporal pRFs and neuronal temporal receptive fields

Our study revealed substantial variability in temporal parameters within an area as well as overlap in temporal parameters across areas (Fig 7A). In particular, areas V1, V2, and V3 contain pRFs with a large range of temporal integration windows that also vary with eccentricity (Fig 9). The temporal overlap between areas (i) may indicate some level of parallel temporal processing across visual areas, and (ii) could be related to the underlying neural architecture as anatomical connections in the visual system are not strictly feedforward from one area to the next due to the many feedback and recurrent connections (Felleman and Van Essen, 1991; Nowak and Bullier, 1997; Lamme and Roelfsema, 2000).

Given this variability and that our temporal estimates are derived from BOLD responses, it is interesting to compare time-to-peak latencies derived from the CST model to those from measurements that afford high temporal resolution (ECoG in humans; single and multiunit recordings in nonhuman primates). While latency estimates from fMRI were more variable than either ECoG or electrophysiology, we strikingly find comparable time-to-peak latencies across measurement modalities. This similarity is despite substantial methodological differences in experimental procedures, number of measurements, stimuli, species, and usage of anesthetics. Across our data, ECoG, and electrophysiology measurements we observed three main similarities (Fig 10). First, earlier visual areas display shorter response latencies than later visual areas. Second, within each area, there is variability in temporal latencies of neurons and voxels, reflecting heterogeneous response properties (Saul and Humphrey, 1990; Henry et al., 2020). Third, across areas, there is a significant degree of overlap of response latencies, suggesting some level of parallel processing in visual cortex (Nowak and Bullier, 1997; Schmolesky et al., 1998).

**Figure 10.**
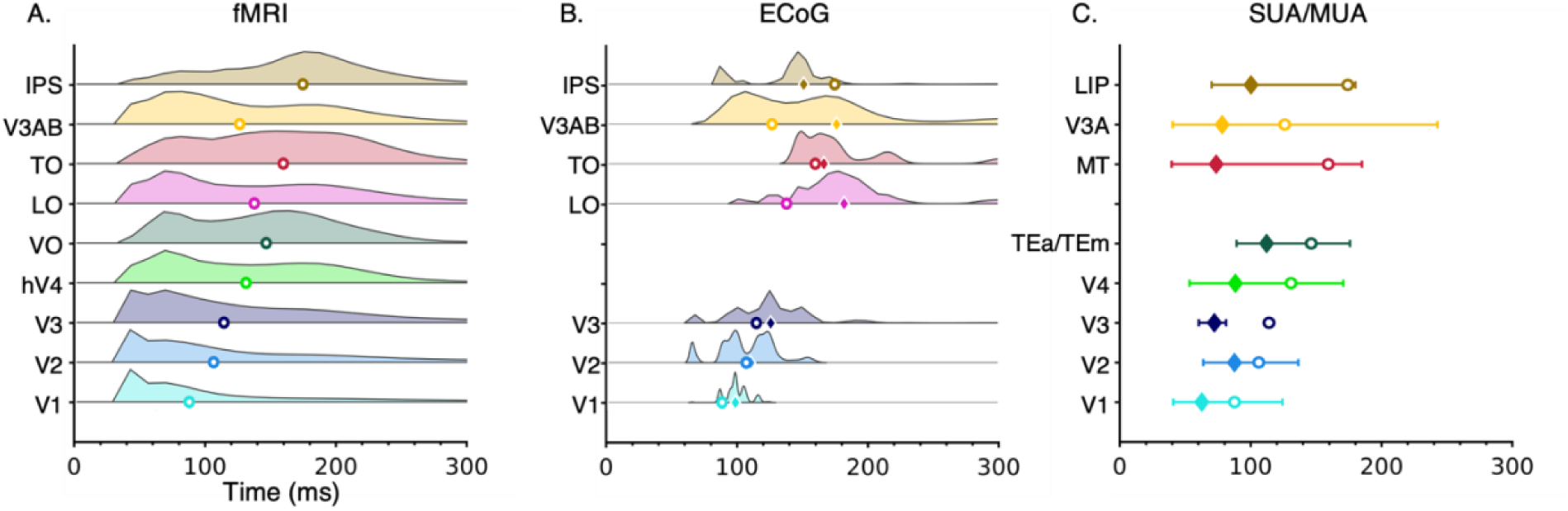
Comparison of estimates of temporal latency in this study to ECoG in humans and electrophysiology in nonhuman primates. In all panels each row illustrates data from a visual area; *x-axis*: time. (A) Distribution of temporal latency estimated from the current study using fMRI. *Open dot*: median of time-to-peak latency. (B) Latency distributions replotted from a human ECoG study (Groen et al., 2022; https://openneuro.org/datasets/ds004194). *Diamond*: median of time-to-peak latency. *Open Circle:* median fMRI latency. (C) Temporal latency ranges from nonhuman primates using electrophysiology; data is aggregated over multiple studies. The central diamonds indicate the average median or mean latency values reported. The whiskers represent the average of minimum/maximum or 10^th^/90^th^ percentile latency values. *Open Circle:* median fMRI latency. Data are from the following studies: *V1* (Raiguel et al., 1989; Knierim and van Essen, 1992; Maunsell and Gibson, 1992; Celebrini et al., 1993; Vogels and Orban, 1994; Nowak et al., 1995; Schmolesky et al., 1998; Bair et al., 2002), *V2* (Raiguel et al., 1989; Nowak et al., 1995; Schmolesky et al., 1998), *V3* (Schmolesky et al., 1998), *V4* (Schmolesky et al., 1998; Chang et al., 2014; Zamarashkina et al., 2020), *TEa/TEm* (Baylis et al., 1987), *MT* (Raiguel et al., 1989, 1999; Schmolesky et al., 1998; Bair et al., 2002; Nakhla et al., 2021), *V3A* (Nakhla et al., 2021), *LIP* (Barash et al., 1991)

Quantitatively, our estimates of time-to-peak latencies in V1-V3 (40-120 ms) are within the range of ECoG: 60 – 80 ms (Martin et al., 2019) to 100 – 120 ms (Zhou et al., 2019; Groen et al., 2022), Fig 10B). Likewise, our estimates of time-to-peak latencies in intermediate (V3AB, LO) and higher-level areas (TO, IPS), ranged from 70 – 200 ms, consistent with ECoG data that ranged from 80 – 150 ms (Martin et al., 2019) to 170 – 200 ms (Groen et al., 2022). However, we note that mean estimates of temporal latencies in humans are consistently slower than in macaques by at least 18 ms (Fig 10C), which may reflect a species difference in temporal processing.

### Hierarchical and parallel processing in visual cortex

Our data reveal that spatiotemporal pRF properties vary across the processing hierarchy as well as between high-level areas in different streams. Consistent with the first hypothesis (Zhou et al., 2018), we found progressive increases in pRF spatial and temporal integration windows as well as in compressive nonlinearities from earlier to later visual areas. This hierarchical structure of spatiotemporal processing of visual information may be achieved via accumulated spatiotemporal pooling across a feedforward neural architecture (Felleman and Van Essen, 1991; Lennie, 1998; Yamins et al., 2014). It is interesting that spatiotemporal characteristics of intermediate visual areas (V3AB, LO, hV4) were similar across streams, but those of higher-level regions (IPS, TO, VO) diverged across streams, with the IPS having the longest temporal integration window and TO having the largest spatial integration window. These findings raise the intriguing hypothesis that the subsequent regions in each of these processing streams would show even more divergence. For example, IPS2-4 in the dorsal stream may have even more different spatiotemporal properties than pFus-faces and mFus-faces in the ventral stream.

### Conclusions

The fMRI techniques based on hemodynamic properties have not been considered as a viable method for studying the timing of human brain activity. However, using a new spatiotemporal pRF model, we demonstrate the possibility of using fMRI to estimate temporal processing at sub-second resolution. This method opens exciting new possibilities for modeling and measuring spatiotemporal processing in the brain, determining the sequential order of neural processing, and examining time-dependent perceptual and cognitive processing beyond the visual system. Examples for future avenues of research include the examination of the spectro-temporal evolution of neural responses in auditory systems (deCharms et al., 1998; Theunissen et al., 2001; Stephens et al., 2013), somatosensory system (Sripati et al., 2006; Reed et al., 2010), visuospatial attention (Kastner et al., 1998), working memory (Zylberberg and Strowbridge, 2017), decision making (Gold and Shadlen, 2007; Louie et al., 2013), and visual capacity (Kupers et al., 2022).

In sum, the spatiotemporal pRF framework offers new avenues to evaluate spatiotemporal processing at the resolution of visual degrees and milliseconds, which can facilitate the understanding of how combined spatial and temporal information is integrated at multiple stages of visual processing.

## Author contributions

I.K. and K.G.S designed research; I.K. and E.R.K performed research; I.K., E.R.K and K.G.S analyzed data; G.L.U contributed to the development of simulation tools; I.K, E.R.K and K.G.S wrote the paper. All authors provided input on the manuscript.

## Acknowledgments

This work was supported by NIH grant R01 EY023915 to KGS. G. L-U. was supported by grants from the Spanish Ministry of Science and Innovation (IJC2020-042887-I; PID2021-123577NA-I00) and Basque Government (PIBA-2022-1-0014). We thank Won Mok Shim for providing resources for pilot data collection. We extend our thanks to Justin Gardner and Brian Wandell for fruitful discussions.

## References

Acerbi L, Ma WJ (2017) Practical Bayesian optimization for model fitting with Bayesian adaptive direct search. Adv Neural Inf Process Syst 30 Available at: https://proceedings.neurips.cc/paper/2017/hash/df0aab058ce179e4f7ab135ed4e641a9-Abstract.html.

Adelson EH, Bergen JR (1985) Spatiotemporal energy models for the perception of motion. J Opt Soc Am A 2:284–299.

Bair W, Cavanaugh JR, Smith MA, Movshon JA (2002) The timing of response onset and offset in macaque visual neurons. J Neurosci 22:3189–3205.

Baldassano C, Chen J, Zadbood A, Pillow JW, Hasson U, Norman KA (2017) Discovering Event Structure in Continuous Narrative Perception and Memory. Neuron 95:709–721.e5.

Barash S, Bracewell RM, Fogassi L, Gnadt JW, Andersen RA (1991) Saccade-related activity in the lateral intraparietal area. I. Temporal properties; comparison with area 7a. J Neurophysiol 66:1095–1108.

Baylis GC, Rolls ET, Leonard CM (1987) Functional subdivisions of the temporal lobe neocortex. J Neurosci 7:330–342.

Brainard DH (1997) The Psychophysics Toolbox. Spat Vis 10:433–436.

Celebrini S, Thorpe S, Trotter Y, Imbert M (1993) Dynamics of orientation coding in area V1 of the awake primate. Vis Neurosci 10:811–825.

Chang M, Xian S, Rubin J, Moore T (2014) Latency of chromatic information in area V4. J Physiol Paris 108:11–17.

Chaudhuri R, Knoblauch K, Gariel M-A, Kennedy H, Wang X-J (2015) A Large-Scale Circuit Mechanism for Hierarchical Dynamical Processing in the Primate Cortex. Neuron 88:419–431.

Conway BR, Livingstone MS (2003) Space-time maps and two-bar interactions of different classes of direction-selective cells in macaque V-1. J Neurophysiol 89:2726–2742.

De Valois RL, Cottaris NP (1998) Inputs to directionally selective simple cells in macaque striate cortex. Proc Natl Acad Sci U S A 95:14488–14493.

De Valois RL, Cottaris NP, Mahon LE, Elfar SD, Wilson JA (2000) Spatial and temporal receptive fields of geniculate and cortical cells and directional selectivity. Vision Res 40:3685–3702.

DeAngelis GC, Ohzawa I, Freeman RD (1993) Spatiotemporal organization of simple-cell receptive fields in the cat’s striate cortex. I. General characteristics and postnatal development. J Neurophysiol 69:1091–1117.

deCharms RC, Blake DT, Merzenich MM (1998) Optimizing sound features for cortical neurons. Science 280:1439–1443.

Dumoulin SO, Wandell BA (2008) Population receptive field estimates in human visual cortex. Neuroimage 39:647–660.

Erhardt EB, Allen EA, Wei Y, Eichele T, Calhoun VD (2012) SimTB, a simulation toolbox for fMRI data under a model of spatiotemporal separability. Neuroimage 59:4160–4167.

Felleman DJ, Van Essen DC (1991) Distributed hierarchical processing in the primate cerebral cortex. Cereb Cortex 1:1–47.

Finzi D, Gomez J, Nordt M, Rezai AA, Poltoratski S, Grill-Spector K (2021) Differential spatial computations in ventral and lateral face-selective regions are scaffolded by structural connections. Nat Commun 12:2278.

Fischl B (2012) FreeSurfer. Neuroimage 62:774–781.

Friston KJ, Fletcher P, Josephs O, Holmes A, Rugg MD, Turner R (1998) Event-related fMRI: characterizing differential responses. Neuroimage 7:30–40.

Gold JI, Shadlen MN (2007) The neural basis of decision making. Annu Rev Neurosci 30:535–574.

Groen IIA, Piantoni G, Montenegro S, Flinker A, Devore S, Devinsky O, Doyle W, Dugan P, Friedman D, Ramsey N, Petridou N, Winawer J (2022) Temporal dynamics of neural responses in human visual cortex. J Neurosci Available at: http://dx.doi.org/10.1523/JNEUROSCI.1812-21.2022.

Handwerker DA, Ollinger JM, D’Esposito M (2004) Variation of BOLD hemodynamic responses across subjects and brain regions and their effects on statistical analyses. Neuroimage 21:1639–1651.

Harvey BM, Dumoulin SO, Fracasso A, Paul JM (2020) A Network of Topographic Maps in Human Association Cortex Hierarchically Transforms Visual Timing-Selective Responses. Curr Biol 30:1424–1434.e6.

Hasson U, Yang E, Vallines I, Heeger DJ, Rubin N (2008) A hierarchy of temporal receptive windows in human cortex. J Neurosci 28:2539–2550.

Hendrikx E, Paul JM, van Ackooij M, van der Stoep N, Harvey BM (2022) Visual timing-tuned responses in human association cortices and response dynamics in early visual cortex. Nat Commun 13:3952.

Henry CA, Jazayeri M, Shapley RM, Hawken MJ (2020) Distinct spatiotemporal mechanisms underlie extra-classical receptive field modulation in macaque V1 microcircuits. Elife 9 Available at: http://dx.doi.org/10.7554/eLife.54264.

Honey CJ, Thesen T, Donner TH, Silbert LJ, Carlson CE, Devinsky O, Doyle WK, Rubin N, Heeger DJ, Hasson U (2012) Slow cortical dynamics and the accumulation of information over long timescales. Neuron 76:423–434.

Hubel DH, Wiesel TN (1968) Receptive fields and functional architecture of monkey striate cortex. J Physiol 195:215–243.

Kastner S, De Weerd P, Desimone R, Ungerleider LG (1998) Mechanisms of directed attention in the human extrastriate cortex as revealed by functional MRI. Science 282:108–111.

Kay KN, Weiner KS, Grill-Spector K (2015) Attention reduces spatial uncertainty in human ventral temporal cortex. Curr Biol 25:595–600.

Kay KN, Winawer J, Mezer A, Wandell BA (2013) Compressive spatial summation in human visual cortex. J Neurophysiol 110:481–494.

Klink PC, Chen X, Vanduffel W, Roelfsema PR (2021) Population receptive fields in nonhuman primates from whole-brain fMRI and large-scale neurophysiology in visual cortex. Elife 10 Available at: http://dx.doi.org/10.7554/eLife.67304.

Knierim JJ, van Essen DC (1992) Neuronal responses to static texture patterns in area V1 of the alert macaque monkey. J Neurophysiol 67:961–980.

Kupers ER, Kim I, Grill-Spector K (2022) A population receptive field modeling framework of sensory suppression in human visual cortex. Available at: https://2022.ccneuro.org/proceedings/0000139.pdf [Accessed April 4, 2023].

Lage-Castellanos A, Valente G, Senden M, De Martino F (2020) Investigating the Reliability of Population Receptive Field Size Estimates Using fMRI. Front Neurosci 14:825.

Lamme VA, Roelfsema PR (2000) The distinct modes of vision offered by feedforward and recurrent processing. Trends Neurosci 23:571–579.

Larsson J, Heeger DJ (2006) Two retinotopic visual areas in human lateral occipital cortex. J Neurosci 26:13128–13142.

Lennie P (1998) Single units and visual cortical organization. Perception 27:889–935.

Lerma-Usabiaga G, Benson N, Winawer J, Wandell BA (2020) A validation framework for neuroimaging software: The case of population receptive fields. PLoS Comput Biol 16:e1007924.

Lewis LD, Setsompop K, Rosen BR, Polimeni JR (2016) Fast fMRI can detect oscillatory neural activity in humans. Proc Natl Acad Sci U S A 113:E6679–E6685.

Lindquist MA, Wager TD (2007) Validity and power in hemodynamic response modeling: a comparison study and a new approach. Hum Brain Mapp 28:764–784.

Liu TT (2016) Noise contributions to the fMRI signal: An overview. Neuroimage 143:141–151.

Louie K, Khaw MW, Glimcher PW (2013) Normalization is a general neural mechanism for context-dependent decision making. Proceedings of the National Academy of Sciences 110:6139–6144.

Martin AB, Yang X, Saalmann YB, Wang L, Shestyuk A, Lin JJ, Parvizi J, Knight RT, Kastner S (2019) Temporal dynamics and response modulation across the human visual system in a spatial attention task: an ECoG study. Journal of Neuroscience 39:333–352.

Maunsell JH, Gibson JR (1992) Visual response latencies in striate cortex of the macaque monkey. J Neurophysiol 68:1332–1344.

McKee SP, Taylor DG (1984) Discrimination of time: comparison of foveal and peripheral sensitivity. J Opt Soc Am A 1:620–627.

McLean J, Palmer LA (1989) Contribution of linear spatiotemporal receptive field structure to velocity selectivity of simple cells in area 17 of cat. Vision Res 29:675–679.

Miller KL, Luh WM, Liu TT, Martinez A, Obata T, Wong EC, Frank LR, Buxton RB (2001) Nonlinear temporal dynamics of the cerebral blood flow response. Hum Brain Mapp 13:1–12.

Mineault PJ, Khawaja FA, Butts DA, Pack CC (2012) Hierarchical processing of complex motion along the primate dorsal visual pathway. Proc Natl Acad Sci U S A 109:E972–80.

Murray JD, Bernacchia A, Freedman DJ, Romo R, Wallis JD, Cai X, Padoa-Schioppa C, Pasternak T, Seo H, Lee D, Wang X-J (2014) A hierarchy of intrinsic timescales across primate cortex. Nat Neurosci 17:1661–1663.

Nakhla N, Korkian Y, Krause MR, Pack CC (2021) Neural Selectivity for Visual Motion in Macaque Area V3A. eNeuro 8 Available at: http://dx.doi.org/10.1523/ENEURO.0383-20.2020.

Nishimoto S, Gallant JL (2011) A three-dimensional spatiotemporal receptive field model explains responses of area MT neurons to naturalistic movies. J Neurosci 31:14551–14564.

Nishimoto S, Vu AT, Naselaris T, Benjamini Y, Yu B, Gallant JL (2011) Reconstructing visual experiences from brain activity evoked by natural movies. Curr Biol 21:1641–1646.

Nowak LG, Bullier J (1997) The timing of information transfer in the visual system. In: Extrastriate Cortex in Primates, pp 205–241. Boston, MA: Springer US.

Nowak LG, Munk MHJ, Girard P, Bullier J (1995) Visual latencies in areas V1 and V2 of the macaque monkey. Vis Neurosci 12:371–384.

Pawar AS, Gepshtein S, Savel’ev S, Albright TD (2019) Mechanisms of Spatiotemporal Selectivity in Cortical Area MT. Neuron 101:514–527.e2.

Pitcher D, Ungerleider LG (2021) Evidence for a Third Visual Pathway Specialized for Social Perception. Trends Cogn Sci 25:100–110.

Polimeni JR, Lewis LD (2021) Imaging faster neural dynamics with fast fMRI: A need for updated models of the hemodynamic response. Prog Neurobiol 207:102174.

Prince JS, Charest I, Kurzawski JW, Pyles JA, Tarr MJ, Kay KN (2022) Improving the accuracy of single-trial fMRI response estimates using GLMsingle. Elife 11 Available at: http://dx.doi.org/10.7554/eLife.77599.

Raiguel SE, Lagae L, Gulyàs B, Orban GA (1989) Response latencies of visual cells in macaque areas V1, V2 and V5. Brain Res 493:155–159.

Raiguel SE, Xiao DK, Marcar VL, Orban GA (1999) Response latency of macaque area MT/V5 neurons and its relationship to stimulus parameters. J Neurophysiol 82:1944–1956.

Reed JL, Qi H-X, Zhou Z, Bernard MR, Burish MJ, Bonds AB, Kaas JH (2010) Response properties of neurons in primary somatosensory cortex of owl monkeys reflect widespread spatiotemporal integration. J Neurophysiol 103:2139–2157.

Saul AB, Humphrey AL (1990) Spatial and temporal response properties of lagged and nonlagged cells in cat lateral geniculate nucleus. J Neurophysiol 64:206–224.

Schmolesky MT, Wang Y, Hanes DP, Thompson KG, Leutgeb S, Schall JD, Leventhal AG (1998) Signal timing across the macaque visual system. J Neurophysiol 79:3272–3278.

Simoncelli EP, Heeger DJ (1998) A model of neuronal responses in visual area MT. Vision Res 38:743–761.

Sripati AP, Yoshioka T, Denchev P, Hsiao SS, Johnson KO (2006) Spatiotemporal receptive fields of peripheral afferents and cortical area 3b and 1 neurons in the primate somatosensory system. J Neurosci 26:2101–2114.

Stephens GJ, Honey CJ, Hasson U (2013) A place for time: the spatiotemporal structure of neural dynamics during natural audition. J Neurophysiol 110:2019–2026.

Stigliani A, Jeska B, Grill-Spector K (2017) Encoding model of temporal processing in human visual cortex. Proc Natl Acad Sci U S A 114:E11047–E11056.

Stigliani A, Jeska B, Grill-Spector K (2019) Differential sustained and transient temporal processing across visual streams. PLoS Comput Biol 15:e1007011.

Theunissen FE, David SV, Singh NC, Hsu A, Vinje WE, Gallant JL (2001) Estimating spatio-temporal receptive fields of auditory and visual neurons from their responses to natural stimuli. Network 12:289–316.

Van Essen DC, Gallant JL (1994) Neural mechanisms of form and motion processing in the primate visual system. Neuron 13:1–10.

Vogels R, Orban GA (1994) Activity of inferior temporal neurons during orientation discrimination with successively presented gratings. J Neurophysiol 71:1428–1451.

Wandell BA, Dumoulin SO, Brewer AA (2009) Visual cortex in humans. Encyclopedia of neuroscience 10:251–257.

Wandell BA, Winawer J (2015) Computational neuroimaging and population receptive fields. Trends Cogn Sci 19:349–357.

Watson AB (1986) Temporal sensitivity. Handbook of perception and human performance 1:1– 43.

Watson AB, Ahumada AJ Jr (1985) Model of human visual-motion sensing. J Opt Soc Am A 2:322–341.

Weiner KS, Grill-Spector K (2013) Neural representations of faces and limbs neighbor in human high-level visual cortex: evidence for a new organization principle. Psychol Res 77:74– 97.

Welvaert M, Rosseel Y (2014) A review of fMRI simulation studies. PLoS One 9:e101953.

Wurm MF, Caramazza A (2022) Two “what” pathways for action and object recognition. Trends Cogn Sci 26:103–116.

Yamins DLK, Hong H, Cadieu CF, Solomon EA, Seibert D, DiCarlo JJ (2014) Performance-optimized hierarchical models predict neural responses in higher visual cortex. Proc Natl Acad Sci U S A 111:8619–8624.

Zamarashkina P, Popovkina DV, Pasupathy A (2020) Timing of response onset and offset in macaque V4: stimulus and task dependence. J Neurophysiol 123:2311–2325.

Zhou J, Benson NC, Kay K, Winawer J (2019) Predicting neuronal dynamics with a delayed gain control model. PLoS Comput Biol 15:e1007484.

Zhou J, Benson NC, Kay KN, Winawer J (2018) Compressive Temporal Summation in Human Visual Cortex. J Neurosci 38:691–709.

Zylberberg J, Strowbridge BW (2017) Mechanisms of Persistent Activity in Cortical Circuits: Possible Neural Substrates for Working Memory. Annu Rev Neurosci 40:603–627.

